# Global topology of brain-wide co-fluctuations links task states, personality, and behavioral symptom dimensions

**DOI:** 10.64898/2026.04.30.722005

**Authors:** Chunyin Siu, Saad Pirzada, Cameron Glick, Richard Betzel, Giovanni Petri, Jeremy R Manning, Leanne Williams, Manish Saggar

## Abstract

Functional connectivity in network neuroscience is traditionally characterized using time-averaged correlations between brain regions. While these summaries capture stable large-scale organization, they do not fully reflect the temporal structure of moment-to-moment interactions. Here, we investigate how the order of interaction used to represent brain dynamics shapes the organization recovered from neural data. We compare three interaction representations of fMRI dynamics: regional activation (node time series), pairwise co-fluctuations (edge time series), and higher-order triplet interactions (triangle time series); within a common topological framework using Mapper from topological data analysis (TDA). Across task and resting-state data, Mapper representations derived from pairwise co-fluctuations more distinctly segregate task conditions than activation-based or higher-order representations. This organization reflects structured coordination patterns beyond activation polarity and is driven by high-amplitude interaction events. Beyond task states, modularity quality computed across all Mapper representations is highest for edge time series and selectively associated with stable individual differences: higher modularity relates to higher conscientiousness and lower internalizing and externalizing symptom dimensions. Together, these findings suggest that behaviorally relevant information is reflected in the topology of moment-to-moment brain interactions. Topological analysis of interaction-level dynamics therefore provides a complementary and interpretable framework for linking large-scale neural coordination to cognition, personality, and mental health.

## 1. Introduction

The brain can be conceptualized as a complex network of neurons that self-organizes into distinct brain regions characterized by relatively homogeneous cytoarchitecture and neural activity patterns. At the macroscopic scale, whole-brain function is often studied by examining coordinated activity among these regions over time. Functional connectivity (FC), typically quantified as the Pearson correlation between regional neural activity time series, is the most widely used approach for summarizing such coordination. While FC has proven valuable for characterizing large-scale functional organization of the brain (Bullmore & Sporns, 2009; Craddock et al., 2013; Rogers et al., 2007; Smith, 2012; Smith et al., 2009), it is inherently a time-averaged measure and therefore does not fully capture transient brain dynamics that may be behaviorally or clinically meaningful.

To overcome this limitation, recent work has introduced *edge time series*, which quantify moment-to-moment interactions between pairs of brain regions by computing the framewise product of their standardized activity signals, also referred to as regional co-fluctuations (Faskowitz et al., 2020; Owen et al., 2021; Zamani Esfahlani et al., 2020) (see Fig. 1d). Important to note that edge time series can be generalized beyond brain region pairs to triplets and groups of more brain regions (Santoro et al., 2024). Unlike traditional FC, edge time series and its higher-dimensional generalizations preserve temporal information and enable the investigation of rapid, transient coordination patterns across the brain. As such, they provide a more fine-grained representation of brain dynamics that may capture task-evoked and spontaneous state transitions. In particular, the edge time series is robust and carries subject-specific information suitable for neural fingerprinting (Sporns et al., 2021).

**Fig. 1:**
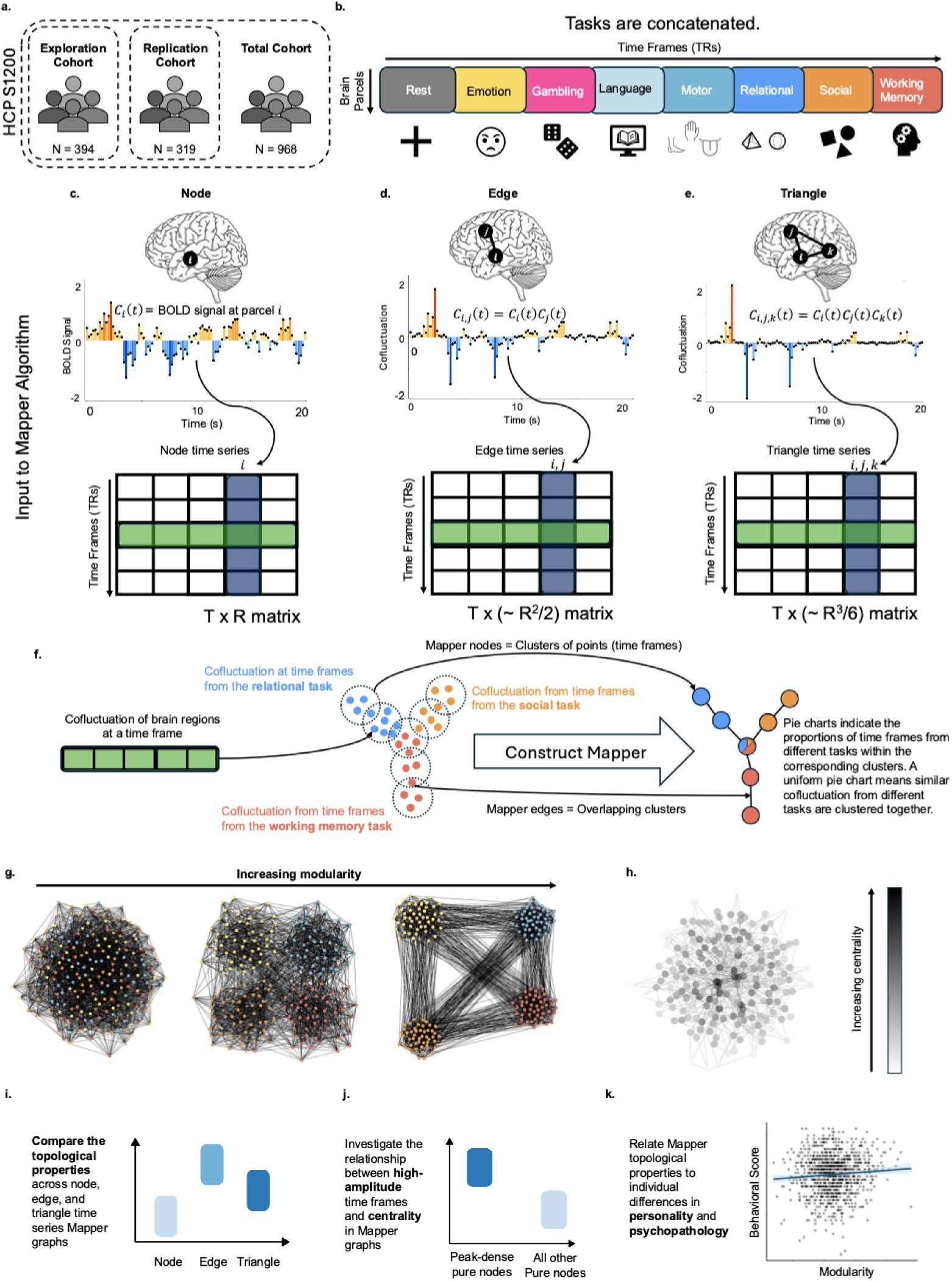
Overview of the approach. **[a]** The Human Connectome Project Young Adult (HCP-YA) dataset was analyzed. After excluding subjects with incomplete data, 968 participants were retained and split into an exploration cohort (N = 394) and an independent replication cohort (N = 319). Behavioral associations were evaluated in the full combined cohort (N = 968). **[b]** For each participant, parcellated BOLD time series were concatenated across resting-state and task fMRI scans. **[c]** Each column of the resulting matrix corresponds to a z-scored regional activation (node) time series for a single brain parcel. **[d]** Edge time series were constructed as the framewise product of pairs of node time series, capturing moment-to-moment co-fluctuations between brain regions. **[e]** Triangle time series were similarly constructed as the framewise product of triplets of node time series, representing higher-order interactions. **[f]** Construction of a Mapper graph from the edge time series for illustration. A similar pipeline was used for node- and triangle time series. Each row of the edge time series matrix (a co-fluctuation vector at a single time frame) is treated as a data point in high-dimensional space. Mapper summarizes the geometry of these data via local clustering, producing a graph whose nodes represent clusters of time frames and whose edges indicate overlapping clusters. Pie charts indicate the proportion of time frames from different tasks within each Mapper node. **[g]** Illustration of graph modularity. Graphs in which nodes of the same type (indicated by color) preferentially cluster together exhibit a higher quality of modularity. **[h]** Illustration of node centrality, with darker shading indicating higher centrality (greater proximity to the average node in the graph). **[i–k]** Schematic of the analysis pipeline, comparing topological properties across Mapper graphs constructed from node, edge, and triangle time series **[i]**, examining the relationship between high-amplitude (peak) time frames and node centrality **[j]**, and relating Mapper-derived topological properties to individual differences in personality and behavioral symptom scales **[k]**.

Analysis of edge time series has revealed the unique role of *high-amplitude frames*, which are brief and intermittent “events” of globally coordinated co-fluctuation (Faskowitz et al., 2020). These infrequent events disproportionately shape time-averaged functional connectivity. Their fundamental importance is underscored by findings that they are constrained by the anatomical connectome (Pope et al., 2021), are conserved across species (Ragone et al., 2024), and synchronize across individuals during shared movie watching (Tanner et al., 2023). High-amplitude frames therefore represent large-scale coordination events.

While edge time series provide a rich description of transient brain dynamics, their extreme high dimensionality poses a fundamental challenge for analysis and interpretation. Conventional dimensionality reduction techniques necessarily compress high-dimensional data into a low-dimensional space, often introducing information loss by collapsing distinct local structures. To address this limitation, we employ Mapper, a tool from topological data analysis that provides a principled approach for embedding high-dimensional data into a low-dimensional representation while preserving essential local relationships (Singh et al., 2007).

Mapper operates by covering the data in a low-dimensional space and performing partial clustering within overlapping regions of this space, resulting in a graph whose nodes represent locally coherent clusters of high-dimensional data points, and whose edges represent overlap of clusters (Singh et al., 2007). Importantly, this construction preserves local distances and neighborhood structure in the original high-dimensional space, while explicitly representing where information loss occurs during dimensionality reduction via node overlap and graph connectivity. In this way, Mapper represents trajectories through high-dimensional space as a network rather than a point-cloud projection, capturing global organization while preserving local geometric structure.

Previous work has demonstrated that Mapper graphs can reveal meaningful structure in neural time series and that graph-derived features can be robustly related to individual differences in behavior and cognition (Geniesse et al., 2022, 2025; Rosenberg et al., 2023; Saggar, Bruno, et al., 2022; Saggar, Shine, et al., 2022; Saggar et al., 2018). In the context of edge time series, the resulting Mapper graph can be interpreted as a low-dimensional summary of the manifold along which brain-wide co-fluctuations evolve over time. To our knowledge, this is among the first applications of Mapper to edge time series data, offering a novel perspective on the temporal organization of brain-wide co-fluctuations.

Here, we investigate whether edge-level dynamics provide a potentially more informative representation of brain state transitions than traditional activation-based (node-level) signals or even higher-order interaction signals (e.g., triangle-level). We compare node-, edge-, and triangle-level representations to determine how interaction order shapes recovered brain organization, and test whether global topological features derived from these representations relate to individual differences in personality and behavioral symptom dimensions. To this end, we construct Mapper graphs from subject-wise edge time series derived from parcellated, concatenated task and resting-state fMRI data from Human Connectome Project (HCP) participants (Barch et al., 2013; Smith et al., 2013).

To quantify task-related organization within these Mapper graphs, we compute the quality of modularity with respect to task labels. Importantly, task labels are not used during Mapper graph construction; rather, they are assigned post hoc to nodes based on the dominant proportion of time frames they contain and are used only to evaluate the degree of segregation across tasks. The quality of modularity is then computed using this predefined partition, without performing community detection on the graph itself. This measure therefore quantifies the extent to which nodes segregate according to task identity (Newman, 2006; Rubinov & Sporns, 2010). We compare the quality of modularity across Mapper graphs constructed from edge time series, traditional node-level activation time series, and triangle-level time series.

Using this framework, we find that Mapper graphs derived from edge time series exhibit structured organization, and they show greater task segregation than graphs derived from node- or triangle-based representations. Moreover, global topological properties of these graphs are associated with individual differences, with the quality of modularity for the edge time series relating to personality traits and self-reported symptom dimensions.

To investigate the origin of the global organization in edge time series Mapper graphs, we analyze the contribution of high-amplitude frames and the mathematical effect of the construction of the edge time series on the distances between activation/co-fluctuation vectors. We identify a prominent role for peak-dense pure nodes, i.e., task-homogenous nodes that are dense with high-amplitude frames, in increasing the quality of modularity, as shuffling their task labels leads to a significant drop in the quality of modularity. We further observe that functional connectivity driven by these nodes corresponds to activation of neurobiologically meaningful, task-specific large-scale brain systems. Our mathematical analysis reveals the quadratic relationship between the node-level distance and edge-level distance between brain (co-) activation vectors. This suggests that the construction of edge time series emphasizes relational structure over absolute activation, making task-specific coordination patterns more coherent in the Mapper graph.

Taken together, these results introduce a framework for visualizing and analyzing dynamic brain interactions and suggest that the global topology of brain-wide co-fluctuations relates to stable behavioral variation.

## 2. Results

We sought to characterize the structure of moment-to-moment brain coordination using representations defined at multiple interaction orders. For each participant, resting-state and task fMRI data were parcellated (Schaefer et al., 2018), z-scored, and concatenated, and each time frame was represented as (i) regional activation patterns (node time series), (ii) pairwise co-fluctuations between regions (edge time series), and (iii) triplet interactions (triangle time series) (Fig. 1c–e). For each representation, we constructed a Mapper graph that groups similar time frames into locally coherent clusters, producing a low-dimensional network representation of brain activity trajectories (Fig. 1f).

Using these parallel representations within a common Mapper framework, we addressed three questions: (a) how interaction order affects task structure, (b) which signal features generate the observed organization, and (c) whether global topological properties relate to stable individual differences. The following sections compare task segregation across representations, examine the role of high-amplitude frames and their anatomical anchors, characterize relationships between node- and edge-level geometry, test associations with personality and behavioral symptom dimensions, and assess reliability across sessions.

### 2.1 The edge time series is better at task profiling than node and triangle time series

We first examined whether different interaction orders provide distinct representations of task structure. Mapper graphs constructed from edge time series exhibited a consistent ring-like organization, in which peripheral nodes were dominated by specific task conditions while centrally located nodes contained mixtures of task time frames (Fig. 2b). For example, resting-state frames preferentially occupied one region of the ring, whereas working memory and social task frames localized to another. This organization was consistently observed across participants (Fig. S3).

**Fig. 2:**
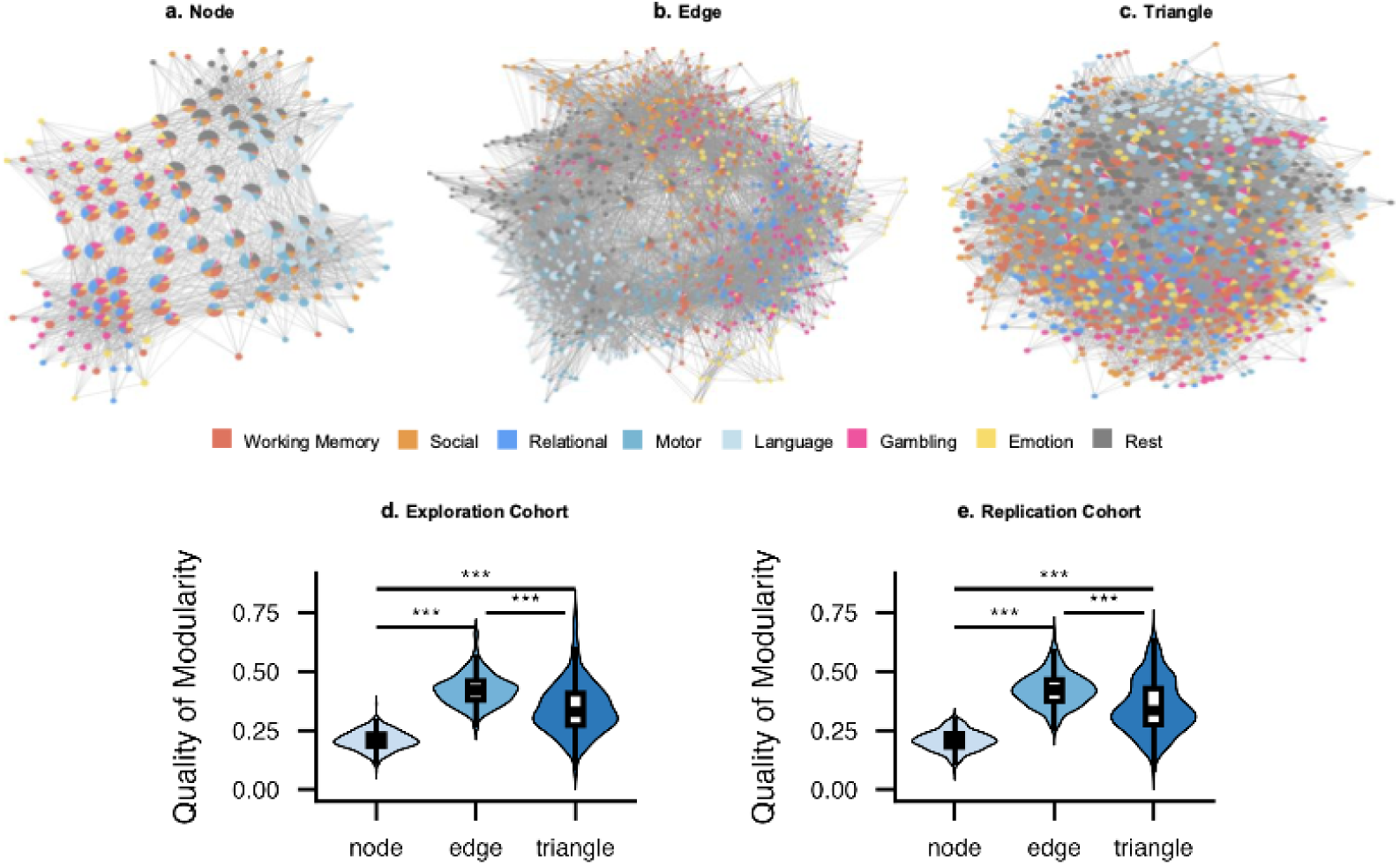
The edge time series is the best at task profiling. **[a – c]** Mapper graphs for the node, edge, and triangle time series of a representative individual. **[d – e]** Distributions of the quality of modularity of edge time series Mapper graphs of the exploration cohort and the replication cohort. See Table S1 for complete statistics. *** means Bonferroni-corrected p-values < 0.001.

In contrast, Mapper graphs derived from node time series showed weak separation between tasks (Fig. 2a). Triangle time series preserved aspects of the ring-like geometry but displayed reduced task specificity compared to edge time series (Fig. 2c).

To quantify task segregation, we computed modularity quality using task identity as a predefined partition of nodes. Each node was assigned a task label based on the dominant proportion of time frames it contained. Across participants in the exploration cohort (N = 394), edge time series showed higher modularity quality than both node and triangle representations (Fig. 2d; paired t-tests, t = 59.0, 17.6, df = 393, Bonferroni-corrected p < 0.001, see Table S1 for complete statistics). This result replicated in an independent cohort (Fig. 2e).

To rule out the effect of scan session and parcellation, we also reproduced this finding in the exploration cohort with a different scan session (RL as opposed to LR), and with a different parcellation (Schaefer 200 instead of Schaefer 100 parcels) (Fig. S1). These results indicate that, as measured by modularity quality, the edge time series shows stronger task segregation than the node and triangle representations across acquisition and parcellation choices.

We also carried out a separate analysis to rule out the effect of simpler graph statistics like the number of Mapper nodes and Mapper edges. When these statistics are controlled, edge time series still attains the highest mean quality of modularity (see Supplementary Materials Section S2).

Because edge time series have substantially higher dimensionality than node time series, we tested whether dimensionality alone accounted for this difference. We projected all representations onto top principal components prior to Mapper construction, yielding comparable feature dimensionality across representations (Fig. S1e – h, Table S1). This procedure reduced the modularity differences, but the differences are still significant (ps < 0.001), indicating that dimensionality contributes to, but does not fully explain, the stronger task segregation observed for edge time series.

These results establish that interaction order influences the degree of task segregation recovered by the Mapper representation.

We next examined which properties of the time series give rise to this organization.

### 2.2 High-amplitude (peak) frames drive task segregation by potentially acting as hubs

Previous works using edge-time series revealed that the brain-wide co-fluctuations occur in intermittent high-amplitude bursts (Faskowitz et al., 2020). These time points, termed high-amplitude or peak frames, correspond to moments in which many edges simultaneously exhibit large co-fluctuations and disproportionately contribute to time-averaged functional connectivity (Zamani Esfahlani et al., 2020). Under naturalistic stimulation, such events align across individuals (Tanner et al., 2023).

We therefore examined whether peak frames contributed to the task segregation observed in Section 2.1. Specifically, we focused on pure Mapper nodes, i.e., nodes in which >75% of time frames belong to a single task, because mixed nodes might act as bridges between tasks and could reduce segregation. The choice of 75% as the purity threshold did not affect the results, as less than 1% of all nodes have purity level strictly between explained in 60% and 100% (see Supplementary Materials Section S6 for details). For the representative participant shown in Fig. 2b, the distribution of peak frames across the Mapper graph is shown in Fig. 3c, and pure nodes with the highest proportion of peak frames (referred to later as peak-dense pure nodes) are highlighted in Fig. 3d.

**Fig. 3.**
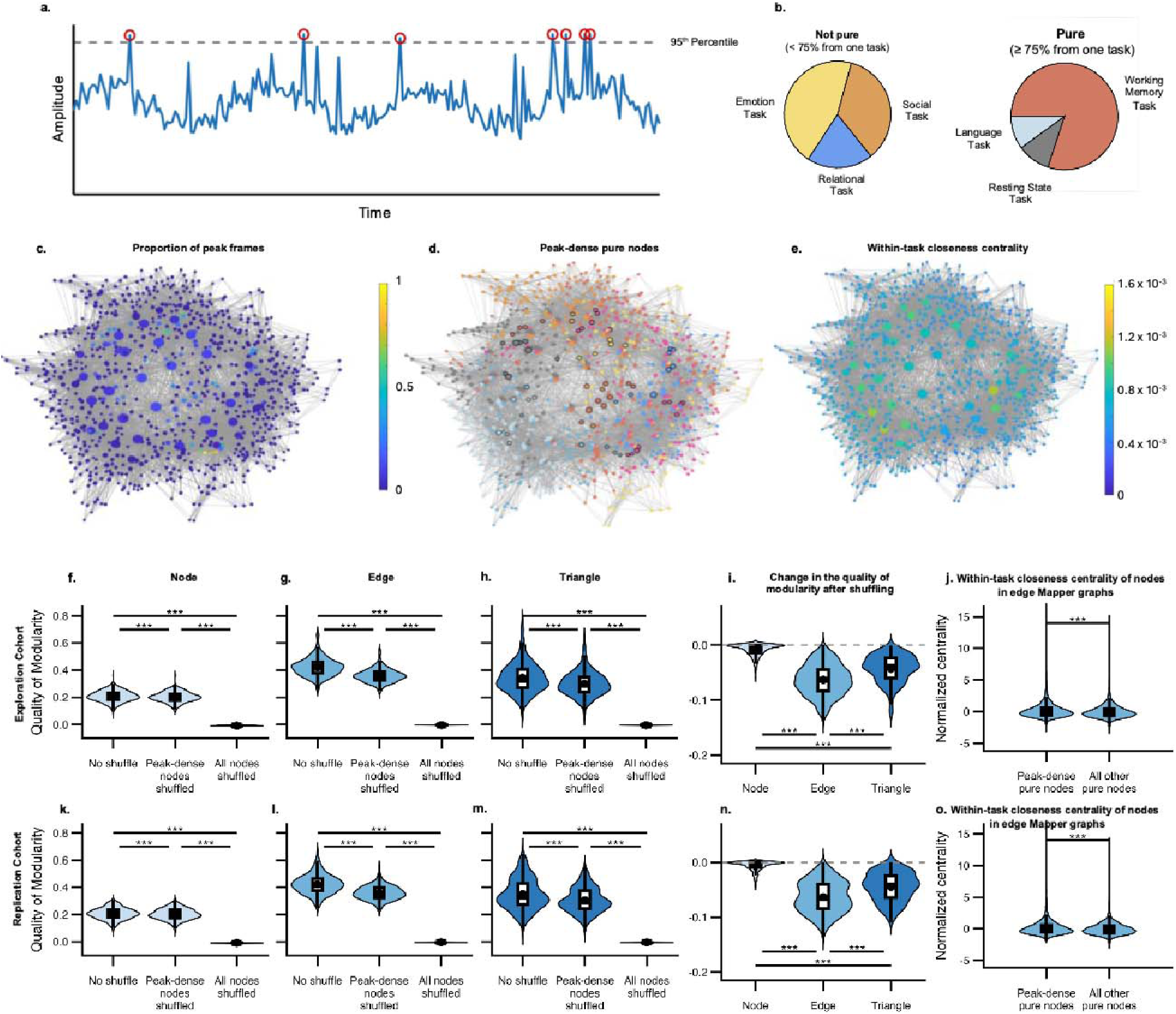
Peak frames drive task segregation as pure peak-dense Mapper nodes serve as task hubs. **[a]** Peak frames are time frames above the 95th percentile in amplitude. **[b]** In a pure Mapper node, over 75% of the time frames in the node belong to the same task. A pure Mapper node is said to be peak-dense if its proportion of peak frames is above the 90th percentile. **[c]** The edge time series Mapper graph in Fig. 2b, colored by the proportion of peak frames. **[d]** The same Mapper graph with peak-dense pure Mapper nodes highlighted. **[e]** The same Mapper graph colored by the within-task centrality. **[f – h]** Distributions (violin plots) of the quality of Mapper modularity with task labels shuffled (not shuffled, only labels of peak-dense pure Mapper nodes shuffled, and all labels shuffled) across simplices (node, edge, triangle) for the exploration cohort. **[i]** Distributions of the change in the quality of modularity upon shuffling task labels of peak-dense pure Mapper nodes across simplices. **[j]** Distributions of within-task centrality of peak-dense pure Mapper nodes and other pure Mapper nodes for the exploration cohort. **[k – o]** Counterparts of Fig. [f – j] for the replication cohort. Throughout all violin plots, pairwise t-tests are done with Bonferroni correction, and asterisks indicate significance (* corrected p < 0.05, ** corrected p < 0.01, *** corrected p < 0.001). See Tables S3 – S5 for complete statistics.

To test peak-dense pure nodes’ contribution to graph organization, we selectively shuffled task labels while preserving graph topology. Four conditions were compared: no shuffling, shuffling peak-dense pure nodes only, shuffling all nodes, and shuffling a matched number of randomly selected nodes. Complete statistics are presented in Table S3. Shuffling only peak-dense pure nodes produced the largest reduction in modularity quality in edge time series graphs, significantly greater than in node or triangle representations. This reduction exceeded that observed when shuffling an equal number of randomly selected nodes, indicating a specific contribution of peak-dense nodes. Similar effects were observed at the individual-subject level (Fig. 3i; Fig. S5). Shuffling all nodes eliminated modularity quality across representations (Fig. 3f–h), showing that the measured modularity reflects alignment between graph structure and task identity.

To understand why these peak-dense nodes exert such influence, we examined their within-task centrality. Peak-dense pure nodes frequently occupied central positions within task-specific clusters (Fig. 3d), suggesting a hub-like role. Thus, we hypothesized that they drive up the quality of modularity by being hubs of their respective tasks, and we tested this hypothesis by measuring the within-task centrality of peak-dense pure nodes. Quantitative analysis confirmed that peak-dense nodes had significantly higher centrality than other pure nodes (t = 20.8, p = 1.43E-94; Fig. 3j, Table S4), and this effect remained significant in a mixed-effects model accounting for subject variability (t = 93.9, p = 0 (too small to be estimated), Table S5).

All findings replicated in an independent cohort (Fig. 3k–o). Controlling for head motion gives similar results (Tables S5 and S7).

Together, these results show that a small subset of high-amplitude frames disproportionately contributes to task segregation in edge time series Mapper graphs.

### 2.3 Task-specific peak frames map onto task-relevant brain systems

Having established that peak-dense pure nodes drive task segregation, we next asked whether these frames correspond to meaningful neurobiological organization.

We first compared functional connectivity (FC) patterns derived from three subsets of time frames: all frames (traditional FC), frames belonging to peak-dense pure nodes, and peak frames alone.

Across tasks, the cohort-wide mean connectivity estimates from all three subsets were highly correlated (cohort-wide mean r > 0.719, lower bound of 95%-confidence interval > 0.712 across connectivity measures and tasks; see Fig. 4a and Table S9). Connectivity derived from peak-dense pure nodes was generally more similar to traditional FC than connectivity derived from peak frames alone. This suggests that the dominant structure of functional connectivity is carried not simply by high-amplitude events, but by high-amplitude events that occur in consistent task-specific configurations.

**Fig. 4:**
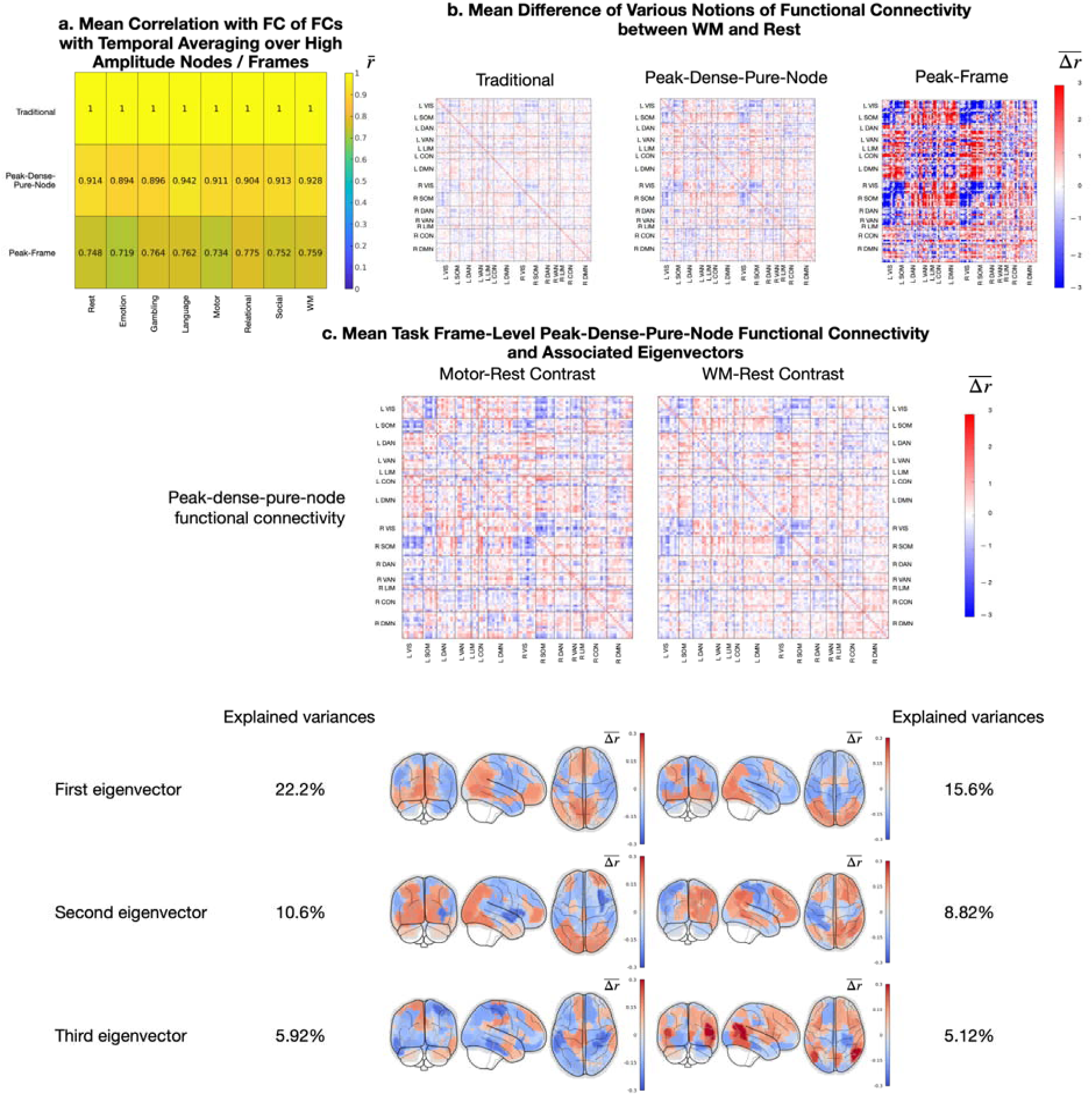
Anchoring task-specific peak frames into task-relevant brain regions. **[a]** Each row is a notion of functional connectivity defined by taking temporal average over a choice of high amplitude nodes or frames, and each column is a task. The colors of the heatmap show the correlation of different forms of functional connectivity with the traditional functional connectivity, averaged across subjects in the exploration cohort. The overall yellowish tone of the heatmap shows the correlations are all close to 1, indicating a linear relationship between different forms of functional connectivity with the traditional functional connectivity. **[b]** The mean difference of functional connectivity matrices between the working memory task and the resting state in the discovery cohort. The more selective we are for high amplitude, the higher the intensity of the heatmap. **[c]** The mean difference in peak-frame functional connectivity matrices from rest for the motor and the working memory tasks and its top 3 eigenvectors (principal components). Note that in the third eigenvector, there are strong (very positive or very negative) loadings in the somatomotor network for the motor task, and strong loadings in the frontoparietal control network for the working memory task.

Consistent with this, the spatial structure of the connectivity matrices was preserved across these estimates (Fig. 4b). Traditional FC exhibited the lowest edge magnitudes, peak-frame FC the highest, and peak-dense-node FC formed an intermediate pattern, suggesting that peak-dense nodes capture the dominant large-scale organization of connectivity while excluding low-amplitude fluctuations.

We next tested whether these connectivity patterns correspond to task-relevant brain systems. Principal components of peak-dense-pure-node connectivity matrices were projected onto the cortical surface (Fig. 4c). During the motor task, a component emphasized the somatomotor network with opposing hemispheric signs consistent with asynchronous limb movements. During the working-memory task, a corresponding component highlighted frontoparietal regions, whereas earlier components reflected general task engagement (e.g., visual involvement). Similar results were observed in the replication cohort (Fig. S9), and approximate equivalence of eigenvectors across scaled matrices is shown in Fig. S8.

Together, these findings indicate that peak-dense frames that organize Mapper topology correspond to structured, task-relevant large-scale brain systems rather than arbitrary fluctuations.

### 2.4 Quadratic Relationship between Node-Level and Edge-Level Distances

To understand why edge time series produce a different graph organization than node time series, we examined how similarity between time frames is defined in each representation. In Mapper, clustering depends on distances between brain states: node time series compare activation vectors, whereas edge time series compare co-fluctuation vectors (Fig. 1c-d).

We begin with a qualitative examination of pairwise distances between time frames across representations. For a representative participant, these distances differed markedly (Fig. 5a). Node-based distances exhibited a checkerboard structure, indicating that some time frames were extremely dissimilar. In contrast, edge-based distances were compressed and lacked these extremes. This suggests that time frames considered opposites in activation space are treated as relatively similar in co-fluctuation space.

**Fig. 5:**
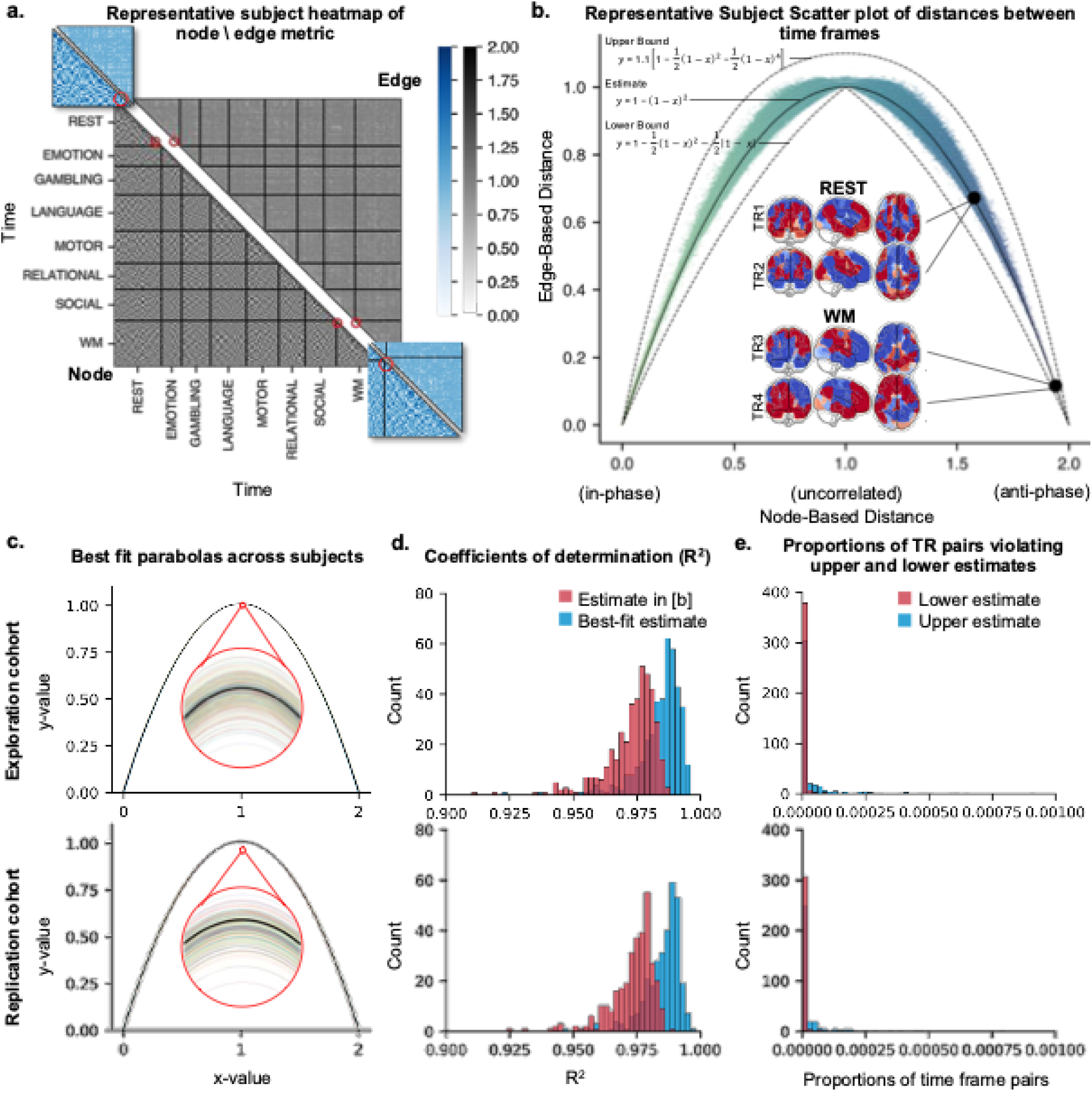
The quadratic relationship between the edge-based distance and node-based distance means that antiphase brain maps are considered similar under the edge-based distance. **[a]** Heatmap of node-based distances (bottom-left) and edge-based distances (top-right) for brain maps at different time frames for a representative subject. The red circles are centered at time frame pairs illustrated in panel [b]. **[b]** Scatter plot of the node-based distances and edge-based distances of time frame pairs for the same subject, with illustrations of the normalized brain maps at time frame pairs furthest apart under the node-based distance (in other words, maximally anti-phase) across tasks and within the resting session. **[c]** The best-fit parabolas for the scatter plots in [b] for all subjects, along with the model parabola *y* = 1 – (1 – *x*)^2^. **[d]** The coefficients of determination of the best-fit parabolas in [c] (blue) and of the model parabola (red). **[e]** The proportion of time frame pairs that violate the upper bound (blue) and the lower bound (red) in [b].

Plotting the two distances against each other revealed a striking nonlinear relationship (Fig. 5b). Across time-frame pairs, the edge-based distance followed a quadratic mapping of the node-based distance:

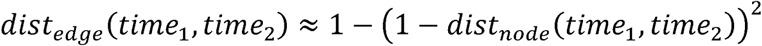

Because correlation distance ranges from 0 (identical) to 2 (anti-phase), this mapping preserves similarity at small distances but compresses large distances, causing anti-correlated activation patterns to appear close in edge space. Thus, edge representations emphasize similarity of interaction structure rather than similarity of activation amplitude.

The approximation in the above quadratic equation is validated in two ways. First, an ordinary least square quadratic approximation is taken for each subject in the exploration cohort, and the best-fit parabolas are all very close to the model parabola *y* = 1 – (1 – *x*)^2^(Fig. 5c), and the coefficients of determination *R*^2^ for the best-fit parabolas and the model parabola are both very close to 1 (cohort-wide minimum *R*^2^ = 91.0%, Fig. 5d). The quadratic approximation can further refined by the upper bound 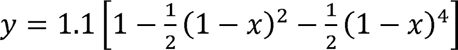 and the lower bound 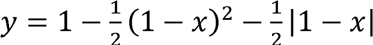, as exemplified by one subject in Fig. 5b. The degree-4 term and the absolute value term are chosen to capture the distribution of points near x = 1, as the lower envelope of the points in Fig. 5b has a sharp kink, and the flatness of the top of the upper envelope suggests the need of a higher power term. The proportion of time point pairs violating either bound is again very low across subjects (cohort-wide maximum proportion of violation = 0.277%, Fig. 5e). These results are replicated in the replication cohort (315 subjects, cohort-wide minimum *R*^2^ = 92.5%, cohort-wide maximum proportion of violation = 0.277%).

For illustration, the most anti-phase activation pairs occurred during the working memory task and during rest (Fig. 5b), yet these pairs remained close in edge space, consistent with the distance transformation.

Together, these findings show that the edge transformation reshapes the geometry of brain states by collapsing oppositely signed activation patterns into nearby points, thereby altering the topology recovered by Mapper. This helps explain the enhanced task segregation observed in Section 2.1: node-based representations treat opposite activation patterns as maximally dissimilar even when they share the same coordination structure, whereas edge representations preserve interaction relationships while discarding polarity. By reducing variability due to global sign and amplitude fluctuations, the representation emphasizes relational structure over absolute activation, making task-specific coordination patterns more coherent in the Mapper graph.

### 2.5 Edge Mapper modularity relates to personality and behavioral symptom scales

We next asked whether the global topological organization of edge time series relates to stable individual differences in behavior. For each participant, the quality of modularity was averaged across runs to reduce measurement variability. We then tested associations between edge modularity (the quality of modularity of an edge time series Mapper graph) and eight behavioral measures, including the five NEO-FFI personality domains, internalizing and externalizing symptom dimensions from the Adult Self-Report (ASR), and fluid intelligence (PMAT).

In the exploration cohort, the quality of modularity showed significant associations with conscientiousness (β = 0.124, uncorrected p = 0.0147), neuroticism (β = 0.112, uncorrected p = 0.0274), internalizing behavior (β = 0.121, uncorrected p = 0.0173), and externalizing behavior (β = -0.157, uncorrected p = 0.00194), and fluid intelligence (β = 0.106, uncorrected p = 0.0375). Among them, the associations with conscientiousness (corrected p = 0.0461), internalizing behavior (corrected p = 0.0461) and externalizing behavior (corrected p = 0.0155) are significant after Benjamini-Hochberg FDR correction. See Table S10 for complete statistics. These three relationships remained significant in after controlling for demographic variables (age and sex), and associations with conscientiousness persisted after accounting for head motion, while the internalizing and externalizing associations were reduced but remained comparable in magnitude (corrected p = 0.0708 and corrected p = 0.0759 respectively, Fig. S11b). See Table S11 for complete statistics for the analysis of the exploration cohort with control variables.

To improve statistical power and reduce behavioral noise, we repeated the analysis in the total cohort using mixed-effects models accounting for family structure, demographics, and head motion. In this analysis, conscientiousness, and internalizing and externalizing symptoms remained significantly associated with the quality of modularity after FDR correction, in the absence of covariate control and when head motion is controlled. When demographics (including family structure) is controlled, the association with externalizing behavior remains significant under FDR correction, those with conscientiousness (p = 0.0635) and internalizing behavior (p = 0.0635) approaches significance under FDR correction. Scatter plots and confidence intervals of regression coefficients are shown in Fig. 6. Complete statistics for the full cohort are presented in Table S12.

**Fig. 6:**
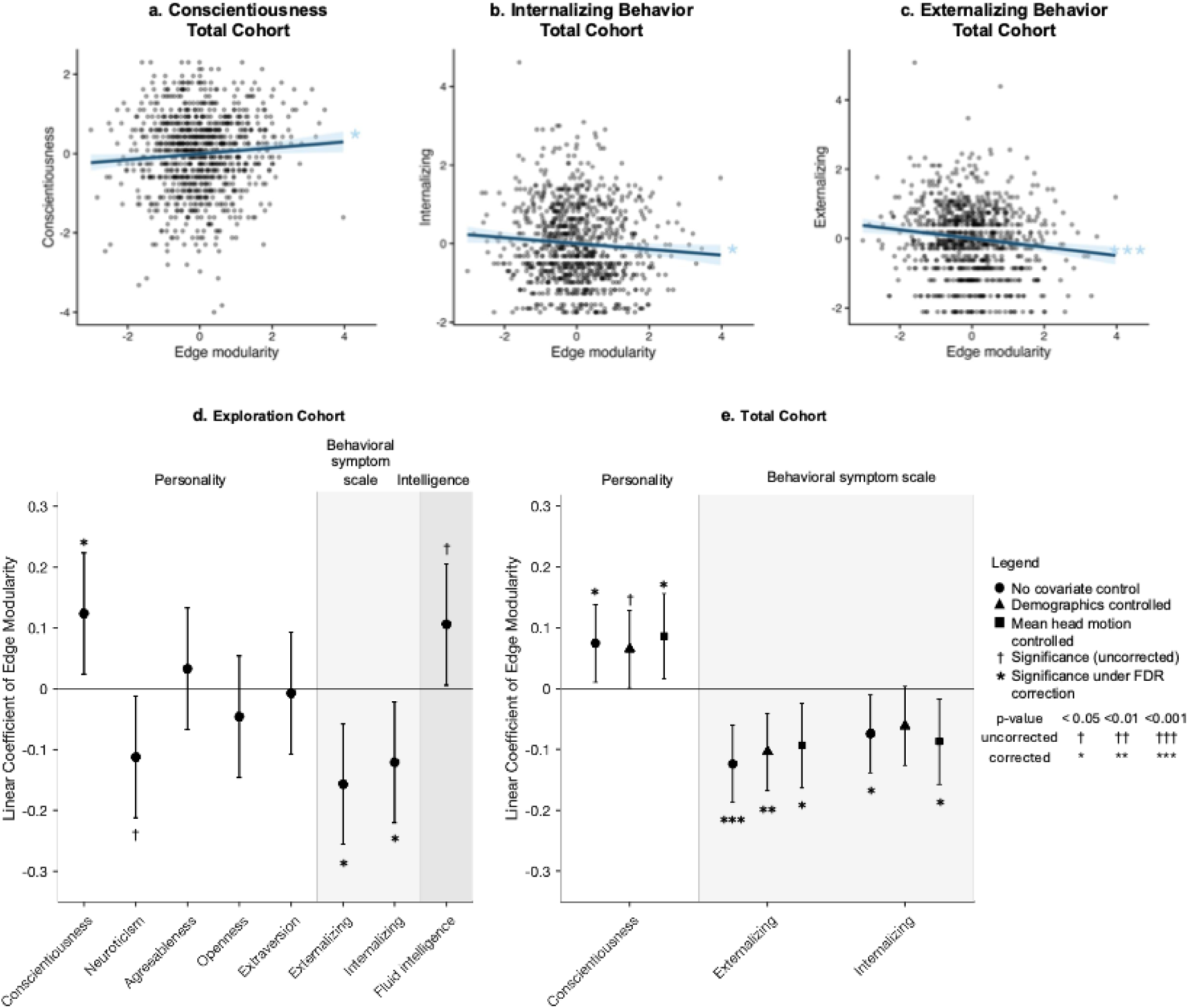
Behavioral correlates of edge modularity (i.e. the quality of modularity for edge time series Mapper graphs). **[a-c]** Edge modularity is significantly correlated with conscientiousness, a domain of temperament in the NEOFFI model, as well as self-reported behavioral symptom scales and fluid intelligence in the total cohort. **[d]** 95% confidence intervals of the linear coefficients of edge modularity in the exploration cohort. **[e]** 95% confidence intervals of the linear coefficients of edge modularity in the total cohort with demographics (age, sex, and family) or head motion controlled.

For comparison, we performed the same analysis using node and triangle time series Mapper graphs (Figs. S10 – S12, Tables S10 – 12). The quality of modularity of node time series Mapper graphs was significantly correlated with agreeableness (without covariate control: β = 0.0734, corrected p = 0.0246, see Fig S12a and Table S12) and fluid intelligence (without covariate control: β = 0.108, corrected p = 0.00179, see Fig S12a and Table S12), but neither correlation remained significant upon controlling for head motion. For triangle time series Mapper graphs, without covariate control, as well as when demographics (including family structure) is controlled, the association of the quality of modularity and externalizing behavior (beta = -0.098, corrected p = 0.0054 (no covariate control) and 0.00264 (demographics controlled)),was significant, but it loses significance upon controlling for head motion (corrected p = 0.115). See Fig S12c and Table S12 for complete statistics.

These findings indicate that global topological organization of moment-to-moment co-fluctuations relates to individual differences in personality and mental health.

## 3. Discussion

Here we asked how the order of interaction used to represent brain activity shapes the latent organization recovered from neural dynamics. By comparing node-, edge-, and triangle-based representations within a common topological framework, we found that pairwise co-fluctuations (edge time series) yield more coherent task-dependent structure than activation-based or higher-order representations. This organization is closely associated with high-amplitude frames and can be understood geometrically through a nonlinear transformation that emphasizes interaction structure over activation polarity. The resulting topological property, i.e., modularity quality, is associated with individual differences in personality and symptom scales and shows stability across sessions. Together, these findings suggest that the topology of dynamic brain interactions provides an interpretable representation linking large-scale coordination patterns to behavior.

### 3.1 Interaction order in large-scale brain dynamics

Complex affective and cognitive phenomena emerge from coordinated activity across distributed brain regions. Traditional node-level analyses describe activity within regions, but they may not capture how relationships between regions structure brain dynamics. This has motivated increasing interest in interaction-based representations, although the appropriate interaction order remains debated. Some studies operationalize interactions using pairwise co-fluctuations such as edge time series and edge-based connectivity (Faskowitz et al., 2020; Zamani Esfahlani et al., 2020), whereas others emphasize triplets or larger collections of regions (Guo et al., 2021; Huang et al., 2017; Owen et al., 2021; Santoro et al., 2024). In parallel, different mathematical frameworks have been proposed to characterize these relationships, including topological approaches such as persistent homology (Petri et al., 2014; Yoon et al., 2024).

By directly comparing node-, edge-, and triangle-level representations within the same analytical framework, we observed that pairwise co-fluctuation structure yielded the most coherent task organization. Rather than indicating that higher-order interactions are uninformative, this finding suggests that interaction order introduces a trade-off between representational expressivity and noise amplification. Pairwise interactions appear to capture dominant coordination patterns while remaining sufficiently constrained to preserve stable structure, whereas higher-order interactions may require stronger regularization or alternative summaries to reveal consistent organization.

In this view, our results reconcile differing perspectives in the literature: large-scale brain dynamics contain higher-order structure, but pairwise interaction representations provide a practical level at which this latent organization becomes detectable in typical neuroimaging data.

### 3.2 High-amplitude frames as task-specific hubs in dynamic brain topology

A growing body of work suggests that functional connectivity is disproportionately shaped by brief high-amplitude events in the edge time series (Ragone et al., 2024; Tanner et al., 2023; Zamani Esfahlani et al., 2020), whereas lower-amplitude frames more closely reflect canonical network structure (Betzel, Cutts, et al., 2023). Previous works revealed that the origins of these high-amplitude frames can be traced to the modular structure of the underlying structural connectome (Pope et al., 2021), and their amplitude is directly related to time-resolved structure-function coupling (Liu et al., 2022). These network dynamics are also linked to endogenous physiological processes, such as hormone fluctuations over the menstrual cycle (Greenwell et al., 2023). Importantly, these co-fluctuation events can be partitioned into discrete states that are stable within individuals (Betzel et al., 2022) and possess a hierarchical organization (Betzel, Cutts, et al., 2023), highlighting their role in shaping person-specific functional brain architecture.

Our results extend this view by showing that these high-amplitude frames are closely linked to the topological organization of brain dynamics. Peak-dense pure nodes exerted a strong influence on the quality of Mapper modularity, and their selective perturbation substantially disrupted task segregation (as shown in Fig. 3). Their elevated within-task centrality further suggests that they act as organizing points around which task-specific brain states cluster.

In prior work using node-based Mapper representations derived from resting-state data, hubs showed broad participation across canonical networks and were interpreted as integrative states shared across ongoing brain dynamics (Saggar, Shine, et al., 2022). The hubs identified here differ in both context and function: during task conditions, high-amplitude interaction patterns form task-specific anchors around which activity organizes. Rather than broadly integrating activity, these hubs stabilize the system within particular coordination regimes. Together, this suggests that dynamic brain organization involves complementary hub types, integrative hubs prominent in unconstrained dynamics and context-specific hubs that anchor coordinated states during goal-directed behavior.

### 3.3 Phase structure revealed by edge representations

We next examined why edge and node representations produce different topological organization. We observed that edge-based distances follow an approximate quadratic transformation of node-based distances (Fig. 6), consistent with prior work suggesting that edge representations reorganize, rather than add to, the information contained in activation patterns (Novelli & Razi, 2022). In particular, we reproduce the observation that the edge time series tends to cluster anticorrelated time frames together (Betzel, Faskowitz, et al., 2023). Under this transformation, activation patterns that are oppositely signed but share the same interaction structure become nearby in edge space. Consequently, Mapper clusters time frames according to coordination structure rather than activation polarity.

This property provides a geometric explanation for the enhanced task segregation observed in edge Mapper graphs. Task-related brain states often involve coordinated but phase-opposed activity across networks, a phenomenon reported in studies of bimanual movement and inter-hemispheric communication (O’Reilly & Elsabbagh, 2021; Wu et al., 2010). By treating antiphase patterns as related states, the edge representation groups fluctuations that belong to the same functional regime while ignoring sign reversals that may arise from oscillatory or context-dependent activity. In this sense, the representation emphasizes phase-consistent coordination patterns, leading to more coherent topological structure.

Together, these findings suggest that edge time series reveal a phase-based organization of brain dynamics that is obscured in activation space, providing a geometric account of their stronger task-dependent topology.

### 3.4 Behavioral relevance of the quality of edge Mapper modularity

Topological properties of brain dynamics have long been linked to behavior, though most studies rely on linear summaries such as functional connectivity. Here, we found that the quality of modularity of the edge time series Mapper graph relates to individual differences in both temperament and symptom scales. This property may reflect the degree to which brain dynamics organize into well-separated coordination regimes over time.

Conscientiousness, a NEO-FFI domain associated with organization, planning, and goal-directed behavior (Costa & McCrae, 1992), showed a positive association with the quality of modularity. Prior work links conscientiousness to structural and functional properties of control-related networks, including lateral prefrontal cortex morphology (DeYoung et al., 2010) and goal-priority systems involving salience and ventral attention networks (Rueter et al., 2018; Sassenberg et al., 2023). In this context, higher quality of modularity may reflect more stable and selective engagement of coordinated neural states, consistent with efficient regulation of attention and behavior.

Conversely, internalizing and externalizing symptoms were associated with reduced quality of modularity. These symptom dimensions have been linked to altered coupling among default mode, frontoparietal, and limbic systems (Kaiser et al., 2015; Thijssen et al., 2021) and to reduced segregation of large-scale networks. Lower quality of modularity in the present framework may therefore indicate less differentiated coordination patterns, consistent with diffuse or interfering neural dynamics.

Together, these relationships suggest a continuum in which well-segregated coordination patterns correspond to organized, goal-directed behavior, whereas reduced segregation corresponds to behavioral dysregulation and inward distress. Rather than serving as a diagnostic marker, the quality of Mapper modularity may index the degree of compartmentalization in ongoing brain dynamics, a property that plausibly supports both behavioral regulation and emotional stability.

### 3.5 Limitations and Future Works

This study has several limitations. First, our findings were derived from the Human Connectome Project dataset, which consists primarily of healthy adults. Although the large sample size allows detection of small brain–behavior associations, replication in independent cohorts and clinical populations will be important to establish generalizability and clarify relevance to psychiatric conditions.

Second, the interpretation of Mapper graphs remains challenging because they summarize very high-dimensional dynamics through a non-linear construction. While we provided geometric and dynamical explanations for the observed structures, the relationship between Mapper topology and underlying neural mechanisms is indirect. Future work could address this by applying the framework to simulated data with known ground-truth dynamics, such as biophysical network models (Deco et al., 2014; Demirtaş et al., 2019; Zhang et al., 2022) or network control theory models (Ceballos et al., 03 2025; Gu et al., 2015), enabling systematic evaluation of what properties of neural systems are recoverable from topological summaries.

Third, our comparison of interaction orders was limited to node-, pairwise-, and triplet-level representations. Higher-order interactions rapidly incur combinatorial growth when defined via time-wise products, making exhaustive evaluation computationally prohibitive. Thus, our results do not imply that higher-order interactions are absent, but rather that pairwise interactions provide a practical level at which stable structure becomes detectable with current data and representations.

Fourth, edge time series were defined using correlation-based co-fluctuations. Alternative definitions of interactions (e.g., nonlinear coupling or model-based interactions) may capture complementary aspects of brain dynamics and could alter the recovered topology. Exploring such formulations will be important for determining which features of neural coordination are most robustly reflected in topological representations.

Finally, our analyses were based on BOLD fMRI signals, which reflect neural activity indirectly through hemodynamic responses. The temporal smoothing imposed by the hemodynamic response function may influence the prominence of high-amplitude events and the apparent phase relationships captured by edge time series. Consequently, the topology described here should be interpreted as characterizing large-scale hemodynamic coordination rather than instantaneous neural interactions. Future work applying the same framework to electrophysiological recordings (e.g., MEG or EEG) will be important to determine which aspects of the observed topology generalize across temporal scales and which are specific to BOLD dynamics.

### 3.6 Conclusions

In summary, we demonstrate that the interaction order used to represent brain activity shapes the organization recovered from neural dynamics. Pairwise co-fluctuations reveal a coherent topological structure that is driven by high-amplitude events, explained by geometric properties of the representation, and associated with stable behavioral variation. These findings position topological analysis as a useful framework for studying large-scale brain coordination and suggest that behaviorally relevant information may be encoded in the global organization of dynamic interactions rather than in regional activation alone.

## 4. Methods

### 4.1 The Human Connectome Project Dataset and Preprocessing

We used the 3T resting state and task-based fMRI data, as well as demographics and behavioral data (sex, age, family, NEOFFI scores, and symptom scale scores) from the de-identified publicly available data from the Human Connectome Project Young Adult (HCP-YA) dataset (Barch et al., 2013; Smith et al., 2013; D. C. Van Essen et al., 2012; David C. Van Essen et al., 2013). As per the HCP protocol guidelines, all participants gave written informed consent for data collection. The HCP scanning protocol was approved by the local Institutional Review Board at Washington University in St. Louis. unrelated subjects who completed the resting state scan and 7 tasks (emotion, gambling, language, motor, relational, social, working memory) were selected to form the <u>exploration cohort</u> of 394 subjects. Another 319 unrelated subjects were selected to form the <u>replication cohort</u>. For behavioral correlation, we repeat the computations in the total cohort of 968 subjects, that comprises both the exploration cohort, the replication cohort, as well as siblings of subjects in both cohorts. For a fairer comparison between resting state activity comparable to task activity, we only include the first 5 minutes of the resting state scan.

For preprocessing, we follow the pipeline described in (Saggar, Shine, et al., 2022), which we briefly describe below. Minimally processed data were first gathered from the HCP database. This minimal processing includes spatial normalization, motion correction, and intensity normalization (Glasser et al., 2013). We additionally processed these data using fMRIPrep 1.5.9 (Esteban et al., 2019).

fMRIPrep anatomical preprocessing included intensity non-uniformity correction with N4BiasFieldCorrection (ANTs 2.2.0), skull-stripping using antsBrainExtraction.sh (OASIS30ANTs template), and tissue segmentation into CSF, white matter, and gray matter using FSL FAST. Nonlinear registration to MNI152 standard spaces was performed with antsRegistration (ANTs 2.2.0).

fMRIPrep functional preprocessing included generating a skull-stripped BOLD reference volume, which was co-registered to the T1w reference using boundary-based registration (FSL FLIRT, 9 degrees of freedom). Head-motion parameters were estimated using MCFLIRT, and BOLD time-series were resampled to native space with motion correction applied. Confound time-series were calculated including framewise displacement (FD), DVARS, and global signals from CSF, white matter, and whole-brain masks.

Temporal masks flagged motion-contaminated frames using FD > 0.5 mm, including one preceding and two following frames. Data were then (i) demeaned and detrended, (ii) regressed against whole-brain, CSF, and white matter signals plus Volterra-expanded motion regressors (excluding censored frames from beta estimation), (iii) interpolated across censored frames using linear estimation, (iv) band-pass filtered (0.009-0.08 Hz), and (v) censored frames removed for final analysis.

### 4.2 Edge Time Series

The node, edge and triangle time series are formed as follows. We use the Schaefer 100 x 7 parcellation scheme (Schaefer et al., 2018). Computationally, the *raw* node time series is a matrix where each entry is the average BOLD signal of the voxels in a parcel (column) at a time frame (row). We obtain the node time series by z-scoring the raw node time series across time parcel by parcel. The edge time series is a matrix where each column corresponds to a pair of parcels, and it is the timewise product of the corresponding columns in the node time series. The triangle time series is similar, with each column corresponding to a triplet of parcels.

To combine different fMRI scans of the same subject, we construct such time series for each scan, and then we concatenate them. In particular, z-scoring is done within each task.

For each instantaneous cofluctuation, which is a row of the edge time series, the amplitude is defined as the ***ℓ_2_*** norm of the row. Mathematically, suppose the parcellation scheme contains ***C_ij_(t)*** parcels. Denote by the cofluctuation of parcel *i* and *j* at time *t*. The instantaneous cofluctuation vector is then ***C(t) = [C_1,2_(t), C_1,3_(t),…,C_m-1,m_(t)].***. The amplitude of ***C(t)*** is

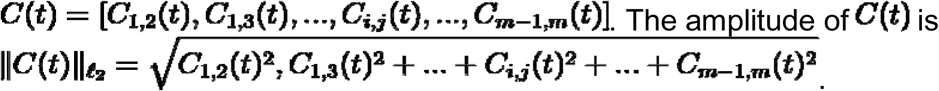

### 4.3 Node-Based Distance and Edge-Based Distance

The node-based distance is simply correlation distance. The edge-based distance is correlation distance for the cofluctuation vectors.

### 4.4 Mapper Graph

The Mapper graph is a graph where nodes correspond to clusters of data points, and edges connect nodes whose corresponding clusters overlap (Singh et al., 2007). Construction of the Mapper graph involves two main steps.

First, filtering: data points are projected into a low-dimensional space using a nonlinear dimensionality reduction method. In this study, we use Isomap, where neighborhood relationships are defined using a k-nearest neighbor (kNN) graph with, where N is the number of data points. This choice balances preservation of local and global structure in the data.

Second, local clustering: the low-dimensional space is partitioned into overlapping bins, and data points within each bin are clustered using a correlation-based distance metric computed from the original dimensional space. The resulting clusters form the nodes of the Mapper graph, and edges are placed between nodes that share data points due to overlap in the binning step.

To reduce sensitivity to manual parameter selection, Mapper parameters were chosen using an adaptive, data-driven strategy (Quah et al., 2025). Specifically, the number of bins (resolution) was determined using kernel density estimation (KDE) to reflect the underlying data density, enabling finer partitioning in regions of higher concentration. The overlap between bins was selected through an iterative procedure to ensure that the largest connected component of the Mapper graph exceeded a predefined threshold (≥ 80% of nodes), promoting a coherent and interpretable topological representation.

A Mapper graph can be annotated using both categorical and quantitative metadata. For quantitative variables (e.g., amplitude), nodes can be colored by the average value within each cluster. For categorical variables (e.g., task identity), nodes can be represented using pie charts indicating the proportion of data points from each category.

### 4.5 Centrality and Modularity

Given a graph (an object made up of nodes and edges), the closeness centrality of a node is the reciprocal of the sum of the distances from the node to all other nodes in the graph (Sabidussi, 1966). The within-task centrality of a node weighs a node by the proportion of time frame in the mode task of the node.

For a graph with a partition of its nodes (grouping of the nodes such that each node belongs to exactly one group), the quality of modularity of the graph with respect to the partition, roughly speaking, measures the discrepancy between the proportions of in-group edges and inter-group edges (Newman, 2006; Rubinov & Sporns, 2010). Precisely, consider a graph with***n_v_***nodes. Denote by ***D_i_*** the number of nodes connected to node ***i***. For every pair of nodes *i* and ***j***, let

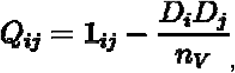

where ***1_ij_***is 1 if nodes and are connected, and it is 0 otherwise. The quality of modularity ***Q*** of the graph with respect to a partition is the sum of the ***Q_ij_***’s for all in-group node pairs ***i, j*** (i.e. ***i*** and ***j*** belong to the same group).

The shuffled quality of modularity of a graph with respect to a collection of nodes is the bootstrap mean of the quality of modularity with the node label (in our case, task labels) randomly shuffled.

### 4.6 Thresholds of Peak Frames and Peak-Dense Nodes

A frame is said to be a peak frame if its amplitude exceeds the 95th percentile. A Mapper node is said to be pure if the proportion of the mode task is at least 75%. A pure Mapper node is said to be a peak-dense pure node if its proportion of peak frames exceeds the 90th percentile. We justify the choice of these thresholds in Supplementary Materials Section S6.

### 4.7 High-Amplitude Functional Connectivity

The peak-frame functional connectivity is the temporal mean of cofluctuation at peak frames. The peak-dense-pure-node functional connectivity is the temporal mean of cofluctuation of frames that belong to peak-dense pure nodes.

### 4.8 FDR correction

FDR correction is applied when the number of tests is large. Generally, Bonferroni correction is done independently for each table in the Supplementary Section. For results regarding personality, behavioral symptoms and fluid intelligence in Section 2.5, within each table (Tables S10 – S12), Benjamini-Hochberg correction is applied for each simplex (node, edge, and triangle) and each covariate control (none, demographics, etc). Further, after the initial analysis in the exploration cohort with no covariate control, features significantly correlated with the quality of modularity after FDR correction are singled out, and only those are tested in the subsequent analyses with covariate control, and subsequently in the total cohort. When no association survive FDR correction in the exploration cohort without covariate control, as in the case of triangle time series, we carry subsequent analysis with features that are significant without correction.

## Data Availability

The data used in this work were originally collected as part of the Human Connectome Project (HCP) (Barch et al., 2013; Smith et al., 2013). We gathered these data directly from the HCP website (https://db.humanconnectome.org).

## Code Availability

Codes to generate Mapper graphs are available at the Mapper repo (https://github.com/braindynamicslab/demapper). Besides codes in our repo, edge time series can also be computed with codes in the edge time series repo (https://github.com/rbetzel/edge-time-series).

## Acknowledgements

Data were provided by the Human Connectome Project, WU-Minn Consortium (Principal Investigators: David Van Essen and Kamil Ugurbil; 1U54MH091657) funded by the 16 NIH Institutes and Centers that support the NIH Blueprint for Neuroscience Research; and by the McDonnell Center for Systems Neuroscience at Washington University.

This research was supported by an MCHRI Faculty Scholar award and an NIH R01 MH127608 to M.S. C.S. was supported by the Croucher Foundation.

## References

Barch, D. M., Burgess, G. C., Harms, M. P., Petersen, S. E., Schlaggar, B. L., Corbetta, M., Glasser, M. F., Curtiss, S., Dixit, S., Feldt, C., Nolan, D., Bryant, E., Hartley, T., Footer, O., Bjork, J. M., Poldrack, R., Smith, S., Johansen-Berg, H., Snyder, A. Z., & Van Essen, D. C. (2013). Function in the human connectome: Task-fMRI and individual differences in behavior. NeuroImage, 80, 169–189.

Betzel, R. F., Cutts, S. A., Greenwell, S., Faskowitz, J., & Sporns, O. (2022). Individualized event structure drives individual differences in whole-brain functional connectivity. NeuroImage, 252(118993), 118993.

Betzel, R. F., Cutts, S. A., Tanner, J., Greenwell, S. A., Varley, T., Faskowitz, J., & Sporns, O. (2023). Hierarchical organization of spontaneous co-fluctuations in densely sampled individuals using fMRI. Network Neuroscience (Cambridge, Mass.), 7(3), 926–949.

Betzel, R. F., Faskowitz, J., & Sporns, O. (2023). Living on the edge: network neuroscience beyond nodes. Trends in Cognitive Sciences, 27(11), 1068–1084.

Bullmore, E., & Sporns, O. (2009). Complex brain networks: graph theoretical analysis of structural and functional systems. Nature Reviews. Neuroscience, 10(3), 186–198.

Ceballos, E. G., Luppi, A. I., Castrillon, G., Saggar, M., Misic, B., & Riedl, V. (03 2025). The control costs of human brain dynamics. Network Neuroscience (Cambridge, Mass.), 9(1), 77–99.

Costa, P. T., & McCrae, R. R. (1992). Revised NEO personality inventory (NEO PI-R) and NEP five-factor inventory (NEO-FFI) : professional manual. Psychological Assessment Resources.

Craddock, R. C., Jbabdi, S., Yan, C.-G., Vogelstein, J. T., Castellanos, F. X., Di Martino, A., Kelly, C., Heberlein, K., Colcombe, S., & Milham, M. P. (2013). Imaging human connectomes at the macroscale. Nature Methods, 10(6), 524–539.

Deco, G., Ponce-Alvarez, A., Hagmann, P., Romani, G. L., Mantini, D., & Corbetta, M. (2014). How Local Excitation–Inhibition Ratio Impacts the Whole Brain Dynamics. The Journal of Neuroscience: The Official Journal of the Society for Neuroscience, 34(23), 7886.

Demirtaş, M., Burt, J. B., Helmer, M., Ji, J. L., Adkinson, B. D., Glasser, M. F., Van Essen, D. C., Sotiropoulos, S. N., Anticevic, A., & Murray, J. D. (2019). Hierarchical heterogeneity across human cortex shapes large-scale neural dynamics. Neuron, 101(6), 1181–1194.e13.

DeYoung, C. G., Hirsh, J. B., Shane, M. S., Papademetris, X., Rajeevan, N., & Gray, J. R. (2010). Testing Predictions From Personality Neuroscience: Brain Structure and the Big Five. Psychological Science, 21(6), 820–828.

Esteban, O., Markiewicz, C. J., Blair, R. W., Moodie, C. A., Isik, A. I., Erramuzpe, A., Kent, J. D., Goncalves, M., DuPre, E., Snyder, M., Oya, H., Ghosh, S. S., Wright, J., Durnez, J., Poldrack, R. A., & Gorgolewski, K. J. (2019). fMRIPrep: a robust preprocessing pipeline for functional MRI. Nature Methods, 16(1), 111–116.

Faskowitz, J., Esfahlani, F. Z., Jo, Y., Sporns, O., & Betzel, R. F. (2020). Edge-centric functional network representations of human cerebral cortex reveal overlapping system-level architecture. Nature Neuroscience, 23(12), 1644–1654.

Geniesse, C., Chowdhury, S., & Saggar, M. (2022). NeuMapper: A scalable computational framework for multiscale exploration of the brain’s dynamical organization. Network Neuroscience (Cambridge, Mass.), 6(2), 467–498.

Geniesse, C., Jahanikia, S., Xie, H., Sonalkar, N. S., Williams, L. M., & Saggar, M. (2025). Topological data analysis reveals rigid brain-state dynamics during self-viewing in trait rumination. In bioRxivorg. bioRxiv. 10.64898/2025.12.27.696696

Glasser, M. F., Sotiropoulos, S. N., Wilson, J. A., Coalson, T. S., Fischl, B., Andersson, J. L., Xu, J., Jbabdi, S., Webster, M., Polimeni, J. R., Van Essen, D. C., & Jenkinson, M. (2013). The minimal preprocessing pipelines for the Human Connectome Project. NeuroImage, 80, 105–124.

Greenwell, S., Faskowitz, J., Pritschet, L., Santander, T., Jacobs, E. G., & Betzel, R. F. (2023). High-amplitude network co-fluctuations linked to variation in hormone concentrations over the menstrual cycle. Network Neuroscience (Cambridge, Mass.), 7(3), 1181–1205.

Gu, S., Pasqualetti, F., Cieslak, M., Telesford, Q. K., Yu, A. B., Kahn, A. E., Medaglia, J. D., Vettel, J. M., Miller, M. B., Grafton, S. T., & Bassett, D. S. (2015). Controllability of structural brain networks. Nature Communications, 6(1), 8414.

Guo, T., Zhang, Y., Xue, Y., Qiao, L., & Shen, D. (2021). Brain Function Network: Higher Order vs. More Discrimination. Frontiers in Neuroscience, 15. 10.3389/fnins.2021.696639

Huang, X., Xu, K., Chu, C., Jiang, T., & Yu, S. (2017). Weak Higher-Order Interactions in Macroscopic Functional Networks of the Resting Brain. The Journal of Neuroscience: The Official Journal of the Society for Neuroscience, 37(43), 10481–10497.

Kaiser, R. H., Andrews-Hanna, J. R., Wager, T. D., & Pizzagalli, D. A. (2015). Large-Scale Network Dysfunction in Major Depressive Disorder: A Meta-analysis of Resting-State Functional Connectivity. JAMA Psychiatry (Chicago, Ill.), 72(6), 603–611.

Liu, Z.-Q., Vázquez-Rodríguez, B., Spreng, R. N., Bernhardt, B. C., Betzel, R. F., & Misic, B. (2022). Time-resolved structure-function coupling in brain networks. Communications Biology, 5(1), 532.

Newman, M. E. J. (2006). Modularity and community structure in networks. Proceedings of the National Academy of Sciences, 103(23), 8577–8582.

Novelli, L., & Razi, A. (2022). A mathematical perspective on edge-centric brain functional connectivity. Nature Communications, 13(1), 2693.

O’Reilly, C., & Elsabbagh, M. (2021). Intracranial recordings reveal ubiquitous in-phase and in-antiphase functional connectivity between homotopic brain regions in humans. Journal of Neuroscience Research, 99(3), 887–897.

Owen, L. L. W., Chang, T. H., & Manning, J. R. (2021). High-level cognition during story listening is reflected in high-order dynamic correlations in neural activity patterns. Nature Communications, 12(1), 5728.

Petri, G., Expert, P., Turkheimer, F., Carhart-Harris, R., Nutt, D., Hellyer, P. J., & Vaccarino, F. (2014). Homological scaffolds of brain functional networks. Journal of the Royal Society, Interface, 11(101), 20140873.

Pope, M., Fukushima, M., Betzel, R. F., & Sporns, O. (2021). Modular origins of high-amplitude cofluctuations in fine-scale functional connectivity dynamics. Proceedings of the National Academy of Sciences of the United States of America, 118(46), e2109380118.

Quah, S. K. L., Madsen, S., Pirzada, S., Jo, B., Uddin, L. Q., Mumford, J. A., Barch, D. M., Gotlib, I. H., Fair, D. A., Poldrack, R. A., & Saggar, M. (2025). Refining RDoC using individual-level task fMRI factor models reveals reproducible brain-wide motifs. In bioRxiv. 10.1101/2025.10.13.682124

Ragone, E., Tanner, J., Jo, Y., Zamani Esfahlani, F., Faskowitz, J., Pope, M., Coletta, L., Gozzi, A., & Betzel, R. (2024). Modular subgraphs in large-scale connectomes underpin spontaneous co-fluctuation events in mouse and human brains. Communications Biology, 7(1), 1–14.

Rogers, B. P., Morgan, V. L., Newton, A. T., & Gore, J. C. (2007). Assessing functional connectivity in the human brain by fMRI. Magnetic Resonance Imaging, 25(10), 1347–1357.

Rosenberg, A. M., Saggar, M., Monzel, A. S., Devine, J., Rogu, P., Limoges, A., Junker, A., Sandi, C., Mosharov, E. V., Dumitriu, D., Anacker, C., & Picard, M. (2023). Brain mitochondrial diversity and network organization predict anxiety-like behavior in male mice. Nature Communications, 14(1), 4726.

Rubinov, M., & Sporns, O. (2010). Complex network measures of brain connectivity: Uses and interpretations. NeuroImage, 52(3), 1059–1069.

Rueter, A. R., Abram, S. V., MacDonald, A. W., Rustichini, A., & DeYoung, C. G. (2018). The goal priority network as a neural substrate of Conscientiousness. Human Brain Mapping, 39(9), 3574–3585.

Sabidussi, G. (1966). The Centrality Index of a Graph. Psychometrika, 31(4), 581–603.

Saggar, M., Bruno, J., Gaillard, C., Claudino, L., & Ernst, M. (2022). Neural resources shift under Methylphenidate: A computational approach to examine anxiety-cognition interplay. NeuroImage, 264(119686), 119686.

Saggar, M., Shine, J. M., Liégeois, R., Dosenbach, N. U. F., & Fair, D. (2022). Precision dynamical mapping using topological data analysis reveals a hub-like transition state at rest. Nature Communications, 13(1), 4791.

Saggar, M., Sporns, O., Gonzalez-Castillo, J., Bandettini, P. A., Carlsson, G., Glover, G., & Reiss, A. L. (2018). Towards a new approach to reveal dynamical organization of the brain using topological data analysis. Nature Communications, 9(1), 1399.

Santoro, A., Battiston, F., Lucas, M., Petri, G., & Amico, E. (2024). Higher-order connectomics of human brain function reveals local topological signatures of task decoding, individual identification, and behavior. Nature Communications, 15(1), 10244.

Sassenberg, T. A., Burton, P. C., Mwilambwe-Tshilobo, L., Jung, R. E., Rustichini, A., Spreng, R. N., & DeYoung, C. G. (2023). Conscientiousness associated with efficiency of the salience/ventral attention network: Replication in three samples using individualized parcellation. NeuroImage, 272, 120081.

Schaefer, A., Kong, R., Gordon, E. M., Laumann, T. O., Zuo, X.-N., Holmes, A. J., Eickhoff, S. B., & Yeo, B. T. T. (2018). Local-Global Parcellation of the Human Cerebral Cortex from Intrinsic Functional Connectivity MRI. Cerebral Cortex (New York, N.Y.: 1991), 28(9), 3095–3114.

Singh, G., Memoli, F., & Carlsson, G. (2007). Topological Methods for the Analysis of High Dimensional Data Sets and 3D Object Recognition. In M. Botsch, R. Pajarola, B. Chen, & M. Zwicker (Eds.), Eurographics Symposium on Point-Based Graphics. The Eurographics Association. 10.2312/SPBG/SPBG07/091-100

Smith, S. M. (2012). The future of FMRI connectivity. NeuroImage, 62(2), 1257–1266.

Smith, S. M., Fox, P. T., Miller, K. L., Glahn, D. C., Fox, P. M., Mackay, C. E., Filippini, N., Watkins, K. E., Toro, R., Laird, A. R., & Beckmann, C. F. (2009). Correspondence of the brain’s functional architecture during activation and rest. Proceedings of the National Academy of Sciences of the United States of America, 106(31), 13040–13045.

Smith, S. M., Vidaurre, D., Beckmann, C. F., Glasser, M. F., Jenkinson, M., Miller, K. L., Nichols, T. E., Robinson, E. C., Salimi-Khorshidi, G., Woolrich, M. W., Barch, D. M., Uğurbil, K., & Van Essen, D. C. (2013). Functional connectomics from resting-state fMRI. Trends in Cognitive Sciences, 17(12), 666–682.

Sporns, O., Faskowitz, J., Teixeira, A. S., Cutts, S. A., & Betzel, R. F. (05 2021). Dynamic expression of brain functional systems disclosed by fine-scale analysis of edge time series. Network Neuroscience (Cambridge, Mass.), 5(2), 405–433.

Tanner, J. C., Faskowitz, J., Byrge, L., Kennedy, D. P., Sporns, O., & Betzel, R. F. (2023). Synchronous high-amplitude co-fluctuations of functional brain networks during movie-watching. Imaging Neuroscience (Cambridge, Mass.), 1, 1–21.

Thijssen, S., Collins, P. F., Weiss, H., & Luciana, M. (2021). The longitudinal association between externalizing behavior and frontoamygdalar resting-state functional connectivity in late adolescence and young adulthood. Journal of Child Psychology and Psychiatry, and Allied Disciplines, 62(7), 857–867.

Van Essen, D. C., Ugurbil, K., Auerbach, E., Barch, D., Behrens, T. E. J., Bucholz, R., Chang, A., Chen, L., Corbetta, M., Curtiss, S. W., Della Penna, S., Feinberg, D., Glasser, M. F., Harel, N., Heath, A. C., Larson-Prior, L., Marcus, D., Michalareas, G., Moeller, S., … Yacoub, E. (2012). The Human Connectome Project: A data acquisition perspective. NeuroImage, 62(4), 2222–2231.

Van Essen, David C., Smith, S. M., Barch, D. M., Behrens, T. E. J., Yacoub, E., & Ugurbil, K. (2013). The WU-Minn Human Connectome Project: An overview. NeuroImage, 80, 62–79.

Wu, T., Wang, L., Hallett, M., Li, K., & Chan, P. (2010). Neural correlates of bimanual anti-phase and in-phase movements in Parkinson’s disease. Brain: A Journal of Neurology, 133(8), 2394–2409.

Yoon, I. H. R., Henselman-Petrusek, G., Yu, Y., Ghrist, R., Smith, S. L., & Giusti, C. (2024). Tracking the topology of neural manifolds across populations. Proceedings of the National Academy of Sciences, 121(46), e2407997121.

Zamani Esfahlani, F., Jo, Y., Faskowitz, J., Byrge, L., Kennedy, D. P., Sporns, O., & Betzel, R. F. (2020). High-amplitude cofluctuations in cortical activity drive functional connectivity. Proceedings of the National Academy of Sciences, 117(45), 28393–28401.

Zhang, M., Sun, Y., & Saggar, M. (2022). Cross-attractor repertoire provides new perspective on structure-function relationship in the brain. NeuroImage, 259(119401), 119401.

