## Supplementary Information for "Global topology of brain-wide co-fluctuations links task states, personality, and behavioral symptom dimensions"

**Supplementary Materials**

Throughout this manuscript, asterisks mean significance after FDR correction: * corrected p < 0.05, ** corrected p < 0.01, *** corrected p < 0.001, “n.s.” corrected p $\geq$

0.05.

**S1 More on Mean Quality of Modularity with Different Parcellations and a Different Cohort**

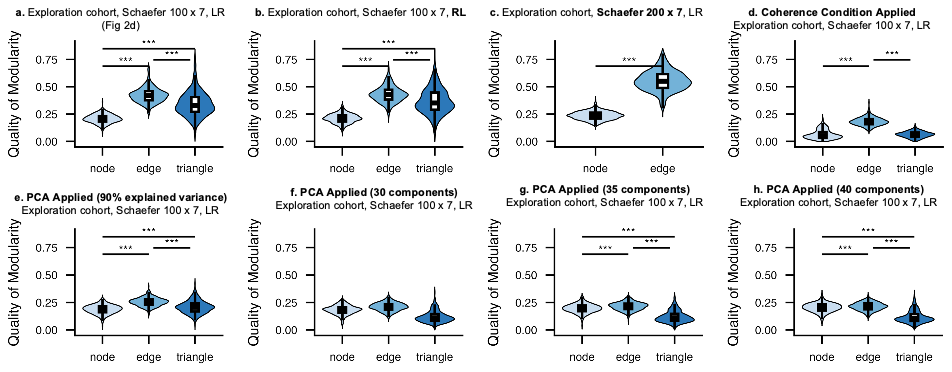

Fig. S1: Analogues of Fig. 1d for different parcellations, cohorts, and data processing methods

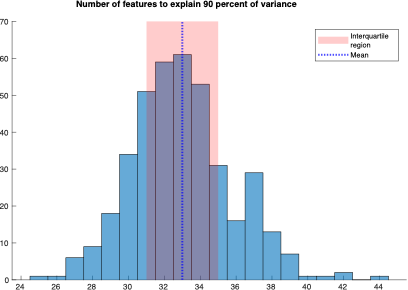

Fig. S2: Distribution of number of principal components to explain 95% variance in the node, edge and triangle time series for the exploration cohort (scan session: LR, parcellation: Schaefer100x7).

We replicate the mean difference of the quality of modularity for the node, edge and triangle time series Mapper graphs in Fig. 1d in the following ways. In the exploration cohort, we replicate with a different scan session (RL as opposed to the LR; Fig. S1b), and with a different parcellation (Schaefer 200 x 7 as opposed to Schaefer 100 x 7; Fig. S1c). Further, we investigate the effect of dimension reduction (Fig. S1e - h) and adding a coherence condition (Fig. S1d) as in (Santoro et al., 2024) on the quality of Mapper modularity. The dimension reduction scheme uses PCA to reduces the dimensions of the edge and triangle time series to either retain the top 90% variance (Fig. S1e) or the top 30, 35, or 40 principal components (Fig. S1f - h). For most people, the 30 – 40 components explain 90% of the variance in the node time series (Fig. S2). By coherence condition (Fig. S1d), we mean we keep only cofluctuation whose absolute z-score is at least 1, and we replace all entries in the node time series with their absolute values, and sign entries in the triangle time series so that they are positive whenever the constituent three values have the same sign, and they are negative if otherwise.

Complete statistics of the comparison of mean quality of modularity are summarized in Table S1.

Table S1: Complete statistics for paired t-tests of mean quality of modularity (Fig 2d-e, S1).

| **Condition** | **Co-hort** | **Ses-sion** | **Parcellation** | **Sim-plex 1** | **Sim-plex 2** | **t-stat** | **DF** | **p-value** | **Cohen's d** | **Sig (Bonferroni corr)** |
| --- | --- | --- | --- | --- | --- | --- | --- | --- | --- | --- |
| Raw Features | one | LR | schaefer100x7 | edge | node | 59 | 393 | 2.5E-197 | 2.97 | *** |
| Raw Features | one | LR | schaefer100x7 | edge | triangle | 17.6 | 393 | 2.41E-51 | 0.884 | *** |
| Raw Features | one | LR | schaefer100x7 | triangle | node | 23.8 | 393 | 2.32E-78 | 1.2 | *** |
| Raw Features | two | LR | schaefer100x7 | edge | node | 52.7 | 318 | 3.86E-159 | 2.95 | *** |
| Raw Features | two | LR | schaefer100x7 | edge | triangle | 14.7 | 318 | 1.28E-37 | 0.822 | *** |
| Raw Features | two | LR | schaefer100x7 | triangle | node | 23.1 | 318 | 4.52E-70 | 1.29 | *** |
| Raw Features | one | RL | schaefer100x7 | edge | node | 59.9 | 396 | 1.16E-200 | 3.01 | *** |
| Raw Features | one | RL | schaefer100x7 | edge | triangle | 13.3 | 395 | 1.14E-33 | 0.669 | *** |
| Raw Features | one | RL | schaefer100x7 | triangle | node | 27.1 | 395 | 6.2E-92 | 1.36 | *** |
| Raw Features | one | LR | schaefer200x7 | edge | node | 67.1 | 392 | 5.17E-217 | 3.39 | *** |
| Coherence | one | LR | schaefer100x7 | edge | node | 42.3 | 386 | 5.17E-147 | 2.15 | *** |
| Coherence | one | LR | schaefer100x7 | edge | triangle | 51.3 | 378 | 2.17E-172 | 2.64 | *** |
| Coherence | one | LR | schaefer100x7 | triangle | node | -0.0551 | 384 | 0.956 | -0.00281 | n.s. |
| PCA (90% variance) | one | LR | schaefer100x7 | edge | node | 28.9 | 393 | 2.53E-99 | 1.46 | *** |
| PCA (90% variance) | one | LR | schaefer100x7 | edge | triangle | 16.5 | 393 | 1.01E-46 | 0.83 | *** |
| PCA (90% variance) | one | LR | schaefer100x7 | triangle | node | 7.12 | 393 | 5.17E-12 | 0.359 | *** |
| PCA (30 comp) | one | LR | schaefer100x7 | edge | node | 10.9 | 390 | 2.93E-24 | 0.55 | *** |
| PCA (30 comp) | one | LR | schaefer100x7 | edge | triangle | 30.1 | 389 | 1.24E-103 | 1.52 | *** |
| PCA (30 comp) | one | LR | schaefer100x7 | triangle | node | -20 | 392 | 5.73E-62 | -1.01 | *** |
| PCA (35 comp) | one | LR | schaefer100x7 | edge | node | 7.3 | 390 | 1.66E-12 | 0.369 | *** |
| PCA (35 comp) | one | LR | schaefer100x7 | edge | triangle | 30.6 | 390 | 1.11E-105 | 1.55 | *** |
| PCA (35 comp) | one | LR | schaefer100x7 | triangle | node | -22.9 | 393 | 2.11E-74 | -1.15 | *** |
| PCA (40 comp) | one | LR | schaefer100x7 | edge | node | 4.38 | 389 | 1.5E-05 | 0.222 | *** |
| PCA (40 comp) | one | LR | schaefer100x7 | edge | triangle | 29.7 | 384 | 1.62E-101 | 1.51 | *** |
| PCA (40 comp) | one | LR | schaefer100x7 | triangle | node | -25.3 | 388 | 5.07E-84 | -1.28 | *** |

**S2 Comparison of Node, Edge, and Triangle Time Series Mapper Graphs with Mapper Graph Size Controlled**

We reproduced the results in Fig. 2d – e with Mapper graph size (i.e. number of Mapper nodes and Mapper edges) controlled. Specifically, we regressed the quality of modularity against two binary variables: whether the Mapper graph is constructed from an edge time series and whether it is constructed from a triangle time series, and two continuous variables: number of nodes and edges in the Mapper graph, as well as one random effect from subjects. In symbols, we have the following formulae:

(1) Quality_of_modularity ~ 1 + is_edge_time_series + is_triangle_time_series + num_Mapper_nodes + num_Mapper_edges + (1 | Subject)

The statistical analytical results from model (1) are summarized in Tables S2. Edge time series mapper graphs still exhibit higher quality of modularity compared with node and triangle time series. For the comparison between the node and triangle time series, there is a reversal in direction (negative coefficients in the Table S2), because of the disproportional increase in complexity of the Mapper graphs for triangle time series.

Table S2: Complete statistics for model (1)

| **Condition** | **Co-hort** | **Ses-sion** | **Parcellation** | **Simplices** | **Coef** | **SE** | **t-stat** | **DF** | **p-value** | **Sig (Bonfe-rroni corr)** |
| --- | --- | --- | --- | --- | --- | --- | --- | --- | --- | --- |
| Raw Features | one | LR | schaefer100x7 | edge-node | 0.0824 | 0.00589 | 14 | 1177 | 2.73E-41 | *** |
| Raw Features | one | LR | schaefer100x7 | triangle-node | -0.0826 | 0.00737 | -11.2 | 1177 | 9.29E-28 | *** |
| Raw Features | one | LR | schaefer100x7 | edge-triangle | 0.165 | 0.00379 | 43.5 | 1177 | 1.99E-247 | *** |
| Raw Features | two | LR | schaefer100x7 | edge-node | 0.0814 | 0.00643 | 12.6 | 952 | 5.22E-34 | *** |
| Raw Features | two | LR | schaefer100x7 | triangle-node | -0.084 | 0.00811 | -10.4 | 952 | 7.05E-24 | *** |
| Raw Features | two | LR | schaefer100x7 | edge-triangle | 0.165 | 0.00422 | 39.2 | 952 | 1.34E-200 | *** |
| Raw Features | one | RL | schaefer100x7 | edge-node | 0.0765 | 0.00661 | 11.6 | 1185 | 2.28E-29 | *** |
| Raw Features | one | RL | schaefer100x7 | triangle-node | -0.094 | 0.00846 | -11.1 | 1185 | 2.43E-27 | *** |
| Raw Features | one | RL | schaefer100x7 | edge-triangle | 0.17 | 0.00431 | 39.6 | 1185 | 4.43E-219 | *** |
| Raw Features | two | RL | schaefer100x7 | edge-node | 0.0876 | 0.00683 | 12.8 | 952 | 8.25E-35 | *** |
| Raw Features | two | RL | schaefer100x7 | triangle-node | -0.0819 | 0.00881 | -9.3 | 952 | 9.4E-20 | *** |
| Raw Features | two | RL | schaefer100x7 | edge-triangle | 0.17 | 0.00443 | 38.3 | 952 | 1.15E-194 | *** |

**S3 More Mapper Graphs from 5 Different Subjects**

We plot Mapper graphs from the subject in Fig. 1 and those from 4 other subjects. These subjects were randomly selected from the exploration cohort.

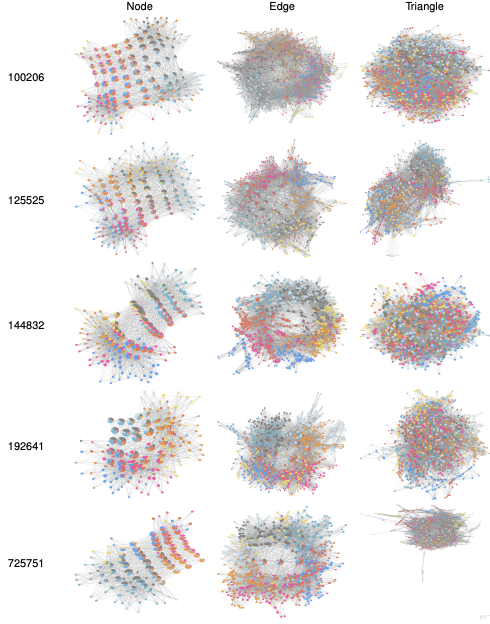

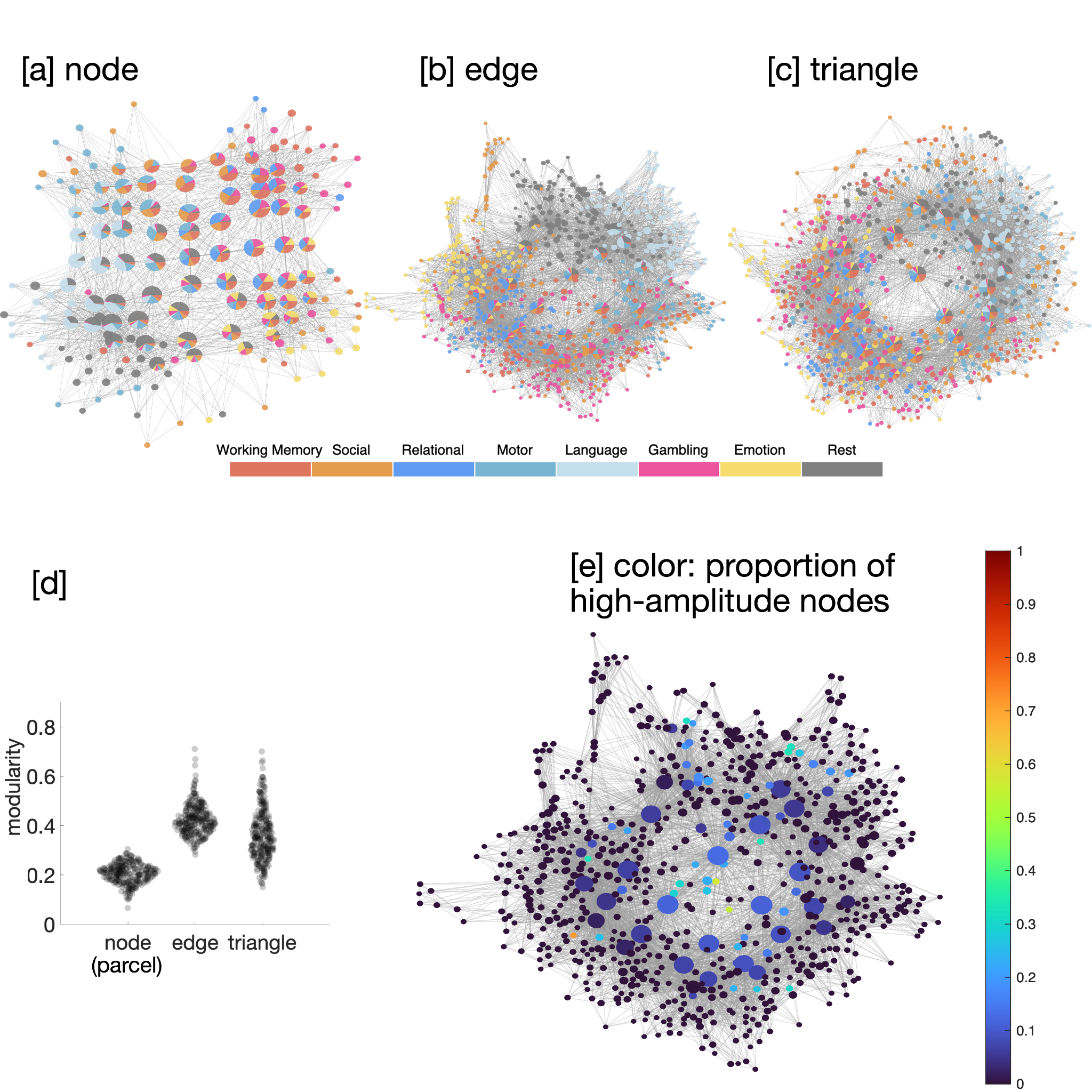

Fig. S3: Mapper graphs for 5 subjects, counterparts of Fig. 1a, 1b and 1c.

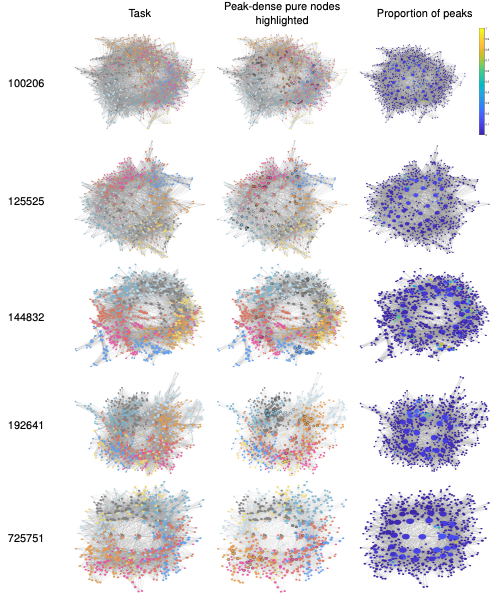

Fig. S4: Mapper graphs for 5 subjects with proportion of peak frames colored and with peak-dense nodes highlighted.

**S4 Distributions of Shuffled Mapper Graphs with the Matched Number of Random Nodes Shuffled**

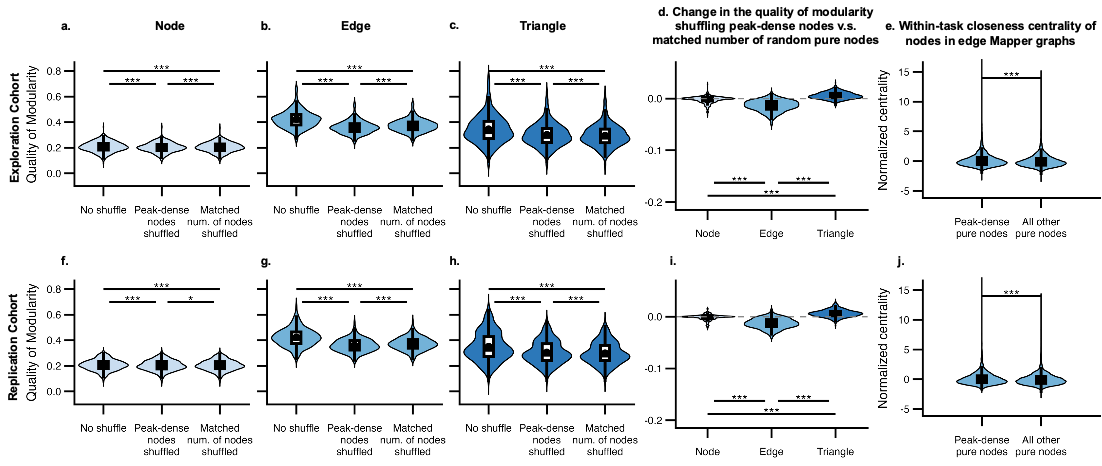

Fig. S5: Distributions of shuffled Mapper graphs; this is the violin plots in Fig. 3 with the column “all nodes shuffled” replaced by “matched number of random nodes shuffled”. Complete statistics in Tables S6 — S7.

Table S3: Complete statistics of the shuffling peak-dense pure nodes experiment (Fig 3)

| **Cohort** | **Session** | **Test Type** | **Simplex** | **t-stat** | **DF** | **p-value** | **Cohen's d** | **Sig (Bonferroni corr)** |
| --- | --- | --- | --- | --- | --- | --- | --- | --- |
| Expl | LR | peak dense minus none | node | -14 | 327 | 2.72E-35 | -0.774 | *** |
| Expl | LR | peak dense minus all | node | 106 | 327 | 4.32E-255 | 5.85 | *** |
| Expl | LR | none minus all | node | 109 | 393 | 1.65E-296 | 5.51 | *** |
| Expl | LR | peak dense minus none | edge | -44.4 | 393 | 2.24E-155 | -2.24 | *** |
| Expl | LR | peak dense minus all | edge | 148 | 393 | 0 | 7.44 | *** |
| Expl | LR | none minus all | edge | 127 | 393 | 2.32E-321 | 6.41 | *** |
| Expl | LR | peak dense minus none | triangle | -34.1 | 393 | 1.53E-119 | -1.72 | *** |
| Expl | LR | peak dense minus all | triangle | 68.1 | 393 | 1.01E-219 | 3.43 | *** |
| Expl | LR | none minus all | triangle | 62.5 | 393 | 3.53E-206 | 3.15 | *** |
| Expl | LR | paired edge vs node | edge-node | -37.6 | 327 | 8.77E-121 | -2.08 | *** |
| Expl | LR | paired edge vs triangle | edge-triangle | -16.8 | 393 | 6E-48 | -0.844 | *** |
| Expl | LR | paired triangle vs node | triangle-node | -25.9 | 327 | 2.95E-81 | -1.43 | *** |
| Repl | LR | peak dense minus none | node | -11.8 | 268 | 2.52E-26 | -0.722 | *** |
| Repl | LR | peak dense minus all | node | 87.9 | 268 | 1.4E-199 | 5.36 | *** |
| Repl | LR | none minus all | node | 94.5 | 318 | 1.01E-234 | 5.29 | *** |
| Repl | LR | peak dense minus none | edge | -39.5 | 318 | 1.34E-124 | -2.21 | *** |
| Repl | LR | peak dense minus all | edge | 127 | 318 | 1.65E-274 | 7.11 | *** |
| Repl | LR | none minus all | edge | 108 | 318 | 4.64E-253 | 6.06 | *** |
| Repl | LR | peak dense minus none | triangle | -30.9 | 318 | 1.14E-97 | -1.73 | *** |
| Repl | LR | peak dense minus all | triangle | 61.9 | 318 | 2.23E-179 | 3.46 | *** |
| Repl | LR | none minus all | triangle | 56.4 | 318 | 1.06E-167 | 3.16 | *** |
| Repl | LR | paired edge vs node | edge-node | -35.8 | 268 | 3.8E-104 | -2.18 | *** |
| Repl | LR | paired edge vs triangle | edge-triangle | -14.2 | 318 | 6.84E-36 | -0.797 | *** |
| Repl | LR | paired triangle vs node | triangle-node | -25.9 | 268 | 7.3E-75 | -1.58 | *** |

Table S4: Complete statistics of t-test for the centrality comparison (Fig 3j, o).

| **Co-hort** | **Ses-sion** | **Mean (peak-dense pure nodes)** | **SD (peak-dense pure nodes)** | **Mean (other pure nodes)** | **SD (other pure nodes)** | **t-stat** | **DF** | **p-value** | **Cohen's d** | **Sig (Bonfe-rroni corr)** |
| --- | --- | --- | --- | --- | --- | --- | --- | --- | --- | --- |
| Expl | LR | 0.0435 | 0.942 | -0.114 | 0.819 | 20.8 | 1.82E+04 | 1.43E-94 | 0.191 | *** |
| Repl | LR | 0.0102 | 0.902 | -0.111 | 0.824 | 15.1 | 1.53E+04 | 6.66E-51 | 0.146 | *** |

Table S5: Complete statistics of centrality comparison with head motion and the subject-level random effect controlled in the model normalized_centrality ~ 1 + is_peak_dense_pure_node + mean_head_motion + (1 | subject) (Fig 3j, o).

| **Cohort** | **Session** | **t-stat** | **DF** | **p-value** | **Coef** | **SE** | **Sig (Bonferroni corr)** |
| --- | --- | --- | --- | --- | --- | --- | --- |
| Expl | LR | 93.9 | 222162 | 0 | 0.244 | 0.0026 | *** |
| Repl | LR | 80.9 | 180945 | 0 | 0.226 | 0.00279 | *** |

Table S6: Complete statistics of the shuffling peak-dense pure nodes experiment with a matched number of random nodes (Fig S5).

| **Cohort** | **Session** | **Test Type** | **Simplex** | **t-stat** | **DF** | **p-value** | **Cohen's d** | **Sig (Bonfe-rroni corr)** |
| --- | --- | --- | --- | --- | --- | --- | --- | --- |
| Expl | LR | peak dense minus none | node | -14 | 327 | 2.72E-35 | -0.774 | *** |
| Expl | LR | peak dense minus matched random | node | -5.77 | 327 | 1.88E-08 | -0.318 | *** |
| Expl | LR | none minus matched random | node | 18.4 | 327 | 1.72E-52 | 1.02 | *** |
| Expl | LR | peak dense minus none | edge | -44.4 | 393 | 2.24E-155 | -2.24 | *** |
| Expl | LR | peak dense minus matched random | edge | -25.5 | 393 | 1.88E-85 | -1.29 | *** |
| Expl | LR | none minus matched random | edge | 40.9 | 393 | 1.5E-143 | 2.06 | *** |
| Expl | LR | peak dense minus none | triangle | -34.1 | 393 | 1.53E-119 | -1.72 | *** |
| Expl | LR | peak dense minus matched random | triangle | 17.1 | 393 | 1.9E-49 | 0.862 | *** |
| Expl | LR | none minus matched random | triangle | 35.4 | 393 | 3.47E-124 | 1.78 | *** |
| Expl | LR | paired edge vs node | edge-node | -19.2 | 327 | 1.13E-55 | -1.06 | *** |
| Expl | LR | paired edge vs triangle | edge-triangle | -32.3 | 393 | 7.93E-113 | -1.63 | *** |
| Expl | LR | paired triangle vs node | triangle-node | 15.2 | 327 | 7.51E-40 | 0.839 | *** |
| Repl | LR | peak dense minus none | node | -11.8 | 268 | 2.52E-26 | -0.722 | *** |
| Repl | LR | peak dense minus matched random | node | -3.46 | 268 | 0.000626 | -0.211 | * |
| Repl | LR | none minus matched random | node | 15.4 | 268 | 1.01E-38 | 0.938 | *** |
| Repl | LR | peak dense minus none | edge | -39.5 | 318 | 1.34E-124 | -2.21 | *** |
| Repl | LR | peak dense minus matched random | edge | -23.8 | 318 | 1.82E-72 | -1.33 | *** |
| Repl | LR | none minus matched random | edge | 34.9 | 318 | 9.48E-111 | 1.95 | *** |
| Repl | LR | peak dense minus none | triangle | -30.9 | 318 | 1.14E-97 | -1.73 | *** |
| Repl | LR | peak dense minus matched random | triangle | 16.1 | 318 | 5.76E-43 | 0.9 | *** |
| Repl | LR | none minus matched random | triangle | 31.8 | 318 | 6.87E-101 | 1.78 | *** |
| Repl | LR | paired edge vs node | edge-node | -19.5 | 268 | 1.95E-53 | -1.19 | *** |
| Repl | LR | paired edge vs triangle | edge-triangle | -29.5 | 318 | 5.08E-93 | -1.65 | *** |
| Repl | LR | paired triangle vs node | triangle-node | 13.9 | 268 | 2.1E-33 | 0.847 | *** |

**S5 Shuffled Mapper Graphs Statistics with Head Motion Controlled**

We repeat all statistical tests in the violin plots in Fig. 3 while controlling for head motion in Table S7.

Table S7: Complete statistics of the shuffling peak-dense pure nodes experiment with head motion controlled.

| **Cohort** | **Session** | **Test Type** | **Simplex** | **Coef** | **SE** | **t-stat** | **DF** | **p-value** | **Sig (Bonfe-rroni corr)** |
| --- | --- | --- | --- | --- | --- | --- | --- | --- | --- |
| Expl | LR | peak dense minus none | node | -0.00844 | 0.000603 | -14 | 326 | 3.44E-35 | *** |
| Expl | LR | peak dense minus all | node | 0.212 | 0.00197 | 108 | 326 | 9.33E-257 | *** |
| Expl | LR | none minus all | node | 0.217 | 0.00194 | 112 | 392 | 1.12E-299 | *** |
| Expl | LR | peak dense minus none | edge | -0.0632 | 0.00138 | -45.8 | 392 | 1.94E-159 | *** |
| Expl | LR | peak dense minus all | edge | 0.364 | 0.00236 | 154 | 392 | 0 | *** |
| Expl | LR | none minus all | edge | 0.427 | 0.00319 | 134 | 392 | 0 | *** |
| Expl | LR | peak dense minus none | triangle | -0.0431 | 0.00118 | -36.7 | 392 | 9.07E-129 | *** |
| Expl | LR | peak dense minus all | triangle | 0.306 | 0.004 | 76.4 | 392 | 1.74E-237 | *** |
| Expl | LR | none minus all | triangle | 0.349 | 0.005 | 69.8 | 392 | 3.1E-223 | *** |
| Expl | LR | paired edge vs node | edge-node | -0.0566 | 0.00147 | -38.4 | 326 | 4.94E-123 | *** |
| Expl | LR | paired edge vs triangle | edge-triangle | -0.0201 | 0.0012 | -16.8 | 392 | 3.58E-48 | *** |
| Expl | LR | paired triangle vs node | triangle-node | -0.0356 | 0.0013 | -27.5 | 326 | 6.53E-87 | *** |
| Repl | LR | peak dense minus none | node | -0.00672 | 0.000569 | -11.8 | 267 | 3.43E-26 | *** |
| Repl | LR | peak dense minus all | node | 0.211 | 0.00236 | 89.6 | 267 | 3.41E-201 | *** |
| Repl | LR | none minus all | node | 0.216 | 0.00222 | 96.9 | 317 | 1.44E-237 | *** |
| Repl | LR | peak dense minus none | edge | -0.0631 | 0.00155 | -40.8 | 317 | 4.02E-128 | *** |
| Repl | LR | peak dense minus all | edge | 0.364 | 0.00269 | 135 | 317 | 1.7E-282 | *** |
| Repl | LR | none minus all | edge | 0.427 | 0.00369 | 116 | 317 | 1.68E-261 | *** |
| Repl | LR | peak dense minus none | triangle | -0.0449 | 0.00139 | -32.4 | 317 | 1.33E-102 | *** |
| Repl | LR | peak dense minus all | triangle | 0.31 | 0.00461 | 67.3 | 317 | 9.02E-190 | *** |
| Repl | LR | none minus all | triangle | 0.355 | 0.00582 | 61.1 | 317 | 1.99E-177 | *** |
| Repl | LR | paired edge vs node | edge-node | -0.0583 | 0.00158 | -36.8 | 267 | 1.48E-106 | *** |
| Repl | LR | paired edge vs triangle | edge-triangle | -0.0181 | 0.00127 | -14.2 | 317 | 8.24E-36 | *** |
| Repl | LR | paired triangle vs node | triangle-node | -0.0399 | 0.00149 | -26.8 | 267 | 1.37E-77 | *** |

**S6 Justification of Choices of Thresholds**

We define a peak frame a one whose amplitude is above the 95th percentile, a pure node as a node whose purity level (proportion of mode task in the node) exceeds 75%, and a pure node to be peak-dense if the proportion of peak frames in the pure node is above the 90th percentile.

Our choice of threshold (95%) for peak frame follows the literature (Zamani Esfahlani et al., 2020).

Our choice of purity level threshold (75%) is immaterial because across cohorts and sessions, less than 1% of the nodes have purity level strictly between 60% and 100% (see first numerical column of Table S8). In fact, over 94% of the nodes have perfect purity (purity level = 100%) (see second numerical column of Table S8). Consequently, any threshold chosen strictly between 60% and 100% results in only a marginal difference in the definition of pure nodes, shifting the proportion from 94% to 95% of all nodes. Further, the distribution of purity is bimodal with a wide trough between the two modes. (Fig. S6) Given the robustness of percentile in general, the choice of the cutoff between the two modes has little impact on the downstream percentile computations.

Table S8: Purity statistics across cohorts and scan sessions.

|  |  | Proportion of nodes with purity level strictly between 60% and 100% | Proportion of perfectly pure nodes (purity level = 100%) |
| --- | --- | --- | --- |
| Exploration cohort | LR | 0.654% | 94.4% |
|  | RL | 0.558% | 94.8% |
| Replication cohort | LR | 0.650% | 94.4% |
|  | RL | 0.553% | 94.9% |

For our choice of threshold for peak-dense nodes, we reproduce Fig. 3f – o for new thresholds 0.8 and 0.95 in place of 0.9 in Fig. S7.

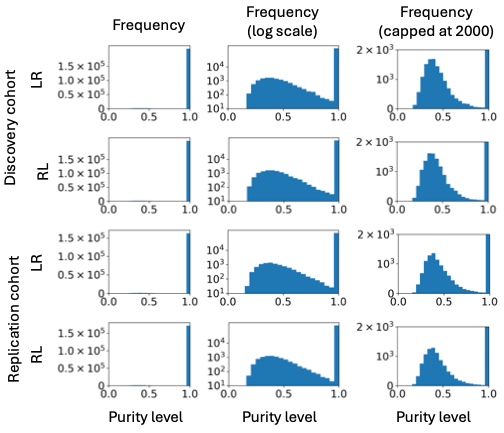

Fig. S6: Histograms of purity level across cohorts and scan sessions.

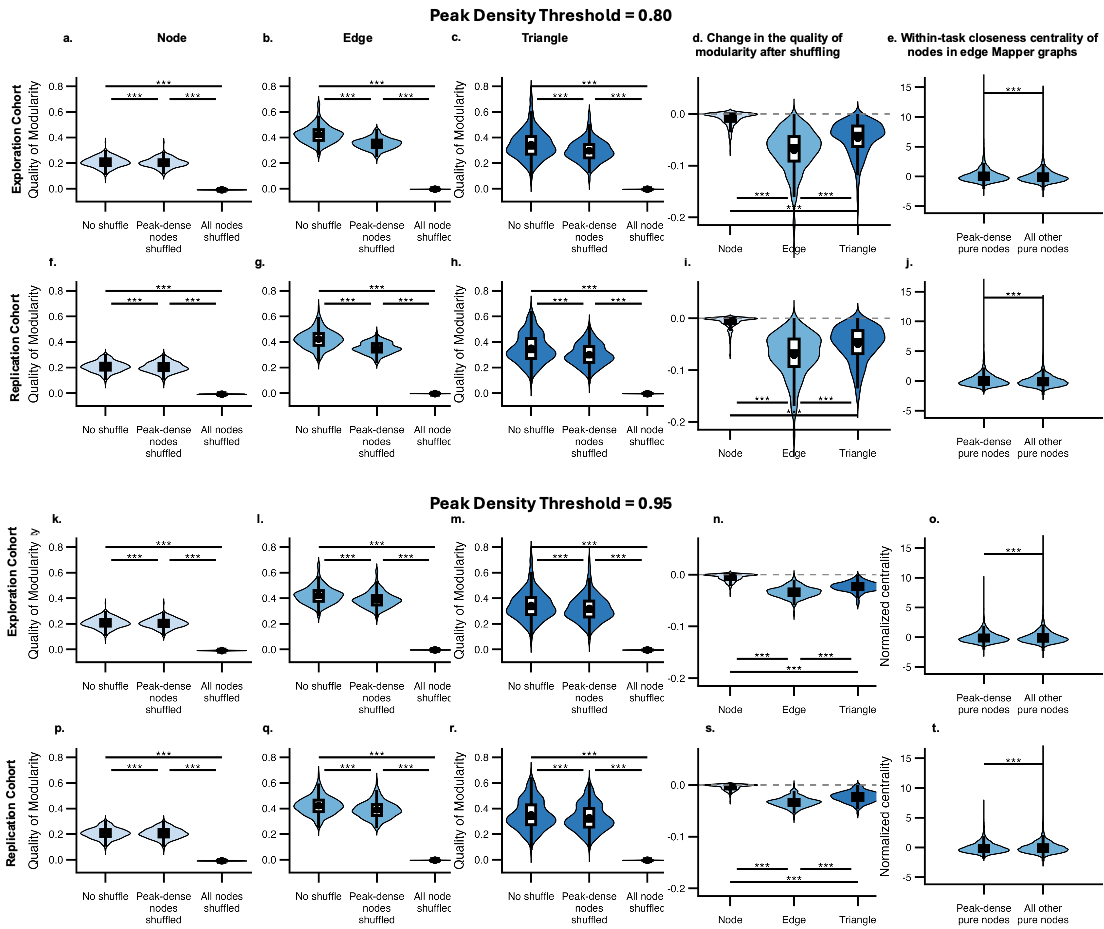
Fig. S7: Analogues of Fig. 3f – o for peak density threshold 0.8 and 0.95.

**S7 More on High-Amplitude Functional Connectivity**

Fig. S8: Eigenvectors of mean difference of task functional connectivity from rest functional connectivity in different notions of functional connectivity defined in terms of peak frames. Note eigenvectors are defined up to sign flip, and eigenvectors in the same columns look alike up to sign.

Table S9: Results of one-sample t-tests on, and the 95%-confidence intervals of, the cohort-wide mean correlation (across region pairs) between the traditional functional connectivity and (1) peak-frame functional connectivity and (2) peak-dense-pure-node functional connectivity. Note that p-values are too small to be estimated.

| **Task** | **Measure** | **N** | **Mean Correlation** | **Confidence Interval (Lower Bound)** | **Confidence Interval (Upper Bound)** | **t-stat** | **p-value** |
| --- | --- | --- | --- | --- | --- | --- | --- |
| **REST** | Peak-Dense-Pure-Node | 394 | 0.914 | 0.906 | 0.923 | 206 | 0 |
| **REST** | Peak-Frame | 394 | 0.748 | 0.742 | 0.754 | 241 | 0 |
| **EMOTION** | Peak-Dense-Pure-Node | 394 | 0.894 | 0.882 | 0.905 | 155 | 0 |
| **EMOTION** | Peak-Frame | 394 | 0.719 | 0.712 | 0.726 | 198 | 0 |
| **GAMBLING** | Peak-Dense-Pure-Node | 394 | 0.896 | 0.886 | 0.907 | 167 | 0 |
| **GAMBLING** | Peak-Frame | 394 | 0.764 | 0.757 | 0.771 | 218 | 0 |
| **LANGUAGE** | Peak-Dense-Pure-Node | 394 | 0.942 | 0.935 | 0.949 | 255 | 0 |
| **LANGUAGE** | Peak-Frame | 394 | 0.762 | 0.756 | 0.769 | 246 | 0 |
| **MOTOR** | Peak-Dense-Pure-Node | 394 | 0.911 | 0.901 | 0.921 | 176 | 0 |
| **MOTOR** | Peak-Frame | 394 | 0.734 | 0.728 | 0.74 | 231 | 0 |
| **RELATIONAL** | Peak-Dense-Pure-Node | 394 | 0.904 | 0.893 | 0.915 | 167 | 0 |
| **RELATIONAL** | Peak-Frame | 394 | 0.775 | 0.768 | 0.782 | 218 | 0 |
| **SOCIAL** | Peak-Dense-Pure-Node | 394 | 0.913 | 0.903 | 0.923 | 183 | 0 |
| **SOCIAL** | Peak-Frame | 394 | 0.752 | 0.745 | 0.759 | 218 | 0 |
| **WM** | Peak-Dense-Pure-Node | 394 | 0.928 | 0.92 | 0.936 | 229 | 0 |
| **WM** | Peak-Frame | 394 | 0.759 | 0.753 | 0.765 | 250 | 0 |

Fig. S9: Analogue of Fig. 4 for the replication cohort.

**S8 More on the Association between the quality of Mapper Modularity and Behavior**

In this section, Benjamini-Hochberg correction is applied for each simplex (node, edge, and triangle) and each covariate control (none, demographics, etc) in each table. Further, after the initial analysis in the exploration cohort with no covariate control, features significantly correlated with the quality of modularity after FDR correction are singled out to be tested in the subsequent analyses with covariate control, and in the total cohort. When no association survive FDR correction in the exploration cohort without covariate control, as in the case of triangle time series, we carry subsequent analysis with features that are significant without correction.

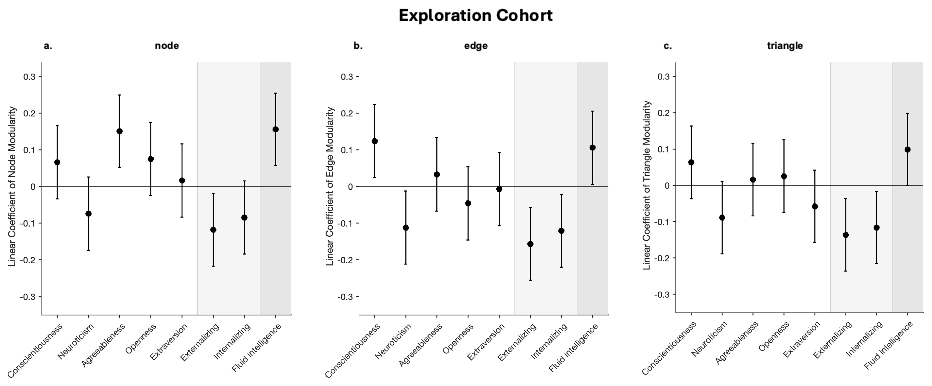

Fig. S10. Two-tailed 95% confidence intervals of the linear coefficients of the quality of Mapper modularity for node, edge and triangle time series in the exploration cohort with no covariate adjustment.

Table S10: Complete statistics of brain-behavior correlation in Fig. S10.

| **Sim-plex** | **Response Variable** | **Control Type** | **Estimate** | **Standard Error** | **DF** | **t-stat** | **p-value** | **Sig** | **p-val (BH corrected)** | **Sig (BH corrected)** |
| --- | --- | --- | --- | --- | --- | --- | --- | --- | --- | --- |
| Node | Conscientiousness | None | 0.0662 | 0.0509 | 382 | 1.3 | 0.194 | n.s. | 0.221 | n.s. |
| Node | Neuroticism | None | -0.074 | 0.0508 | 382 | -1.46 | 0.146 | n.s. | 0.194 | n.s. |
| Node | Agreeableness | None | 0.151 | 0.0504 | 382 | 2.99 | 0.00295 | ** | 0.0118 | * |
| Node | Openness | None | 0.0754 | 0.0508 | 382 | 1.48 | 0.139 | n.s. | 0.194 | n.s. |
| Node | Extraversion | None | 0.0164 | 0.051 | 382 | 0.323 | 0.747 | n.s. | 0.747 | n.s. |
| Node | Externalizing | None | -0.118 | 0.0506 | 382 | -2.33 | 0.0201 | * | 0.0536 | n.s. |
| Node | Internalizing | None | -0.0848 | 0.0508 | 382 | -1.67 | 0.0958 | n.s. | 0.1916 | n.s. |
| Node | Fluid Intelligence | None | 0.156 | 0.0503 | 382 | 3.1 | 0.00211 | ** | 0.0118 | * |
| Edge | Conscientiousness | None | 0.124 | 0.0506 | 382 | 2.45 | 0.0147 | * | 0.046 | * |
| Edge | Neuroticism | None | -0.112 | 0.0507 | 382 | -2.21 | 0.0274 | * | 0.0548 | n.s. |
| Edge | Agreeableness | None | 0.0327 | 0.051 | 382 | 0.643 | 0.521 | n.s. | 0.595 | n.s. |
| Edge | Openness | None | -0.0461 | 0.0509 | 382 | -0.905 | 0.366 | n.s. | 0.488 | n.s. |
| Edge | Extraversion | None | -0.00725 | 0.051 | 382 | -0.142 | 0.887 | n.s. | 0.887 | n.s. |
| Edge | Externalizing | None | -0.157 | 0.0503 | 382 | -3.12 | 0.00194 | ** | 0.01552 | * |
| Edge | Internalizing | None | -0.121 | 0.0506 | 382 | -2.39 | 0.0173 | * | 0.0461 | * |
| Edge | Fluid Intelligence | None | 0.106 | 0.0507 | 382 | 2.09 | 0.0375 | * | 0.06 | n.s. |
| Triangle | Conscientiousness | None | 0.0632 | 0.051 | 381 | 1.24 | 0.216 | n.s. | 0.341 | n.s. |
| Triangle | Neuroticism | None | -0.0887 | 0.0508 | 381 | -1.75 | 0.0818 | n.s. | 0.1636 | n.s. |
| Triangle | Agreeableness | None | 0.0156 | 0.0509 | 381 | 0.306 | 0.76 | n.s. | 0.76 | n.s. |
| Triangle | Openness | None | 0.0251 | 0.0511 | 381 | 0.491 | 0.623 | n.s. | 0.712 | n.s. |
| Triangle | Extraversion | None | -0.058 | 0.051 | 381 | -1.14 | 0.256 | n.s. | 0.341 | n.s. |
| Triangle | Externalizing | None | -0.136 | 0.0506 | 381 | -2.69 | 0.0074 | ** | 0.0592 | n.s. |
| Triangle | Internalizing | None | -0.116 | 0.0506 | 381 | -2.3 | 0.0218 | * | 0.0872 | n.s. |
| Triangle | Fluid Intelligence | None | 0.0989 | 0.0506 | 381 | 1.95 | 0.0514 | n.s. | 0.137 | n.s. |

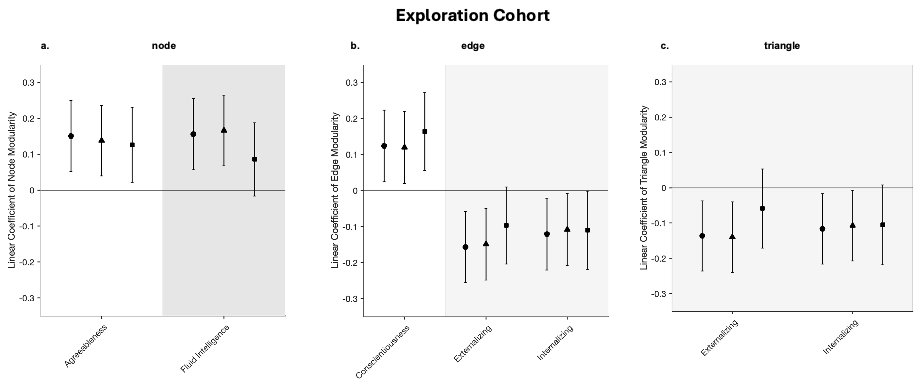

Fig. S11. 95% confidence intervals of the linear coefficients of the quality of Mapper modularity for node, edge and triangle time series in the exploration cohort with demographics (age, sex, and family) and / or head motion controlled.

Table S11: Complete statistics of brain-behavior correlation in Fig. S11.

| **Simplex** | **Response Variable** | **Control Type** | **Estimate** | **Standard Error** | **DF** | **t-stat** | **p-value** | **Sig** | **p-val (BH corrected)** | **Sig (BH corrected)** |
| --- | --- | --- | --- | --- | --- | --- | --- | --- | --- | --- |
| Node | Agreeableness | None | 0.151 | 0.0504 | 382 | 2.99 | 0.00295 | ** | 0.00295 | ** |
| Node | Fluid Intelligence | None | 0.156 | 0.0503 | 382 | 3.1 | 0.00211 | ** | 0.00295 | ** |
| Node | Agreeableness | Demographics | 0.138 | 0.0499 | 380 | 2.77 | 0.00592 | ** | 0.00592 | ** |
| Node | Fluid Intelligence | Demographics | 0.166 | 0.0501 | 380 | 3.31 | 0.00101 | ** | 0.00202 | ** |
| Node | Agreeableness | Head Motion | 0.126 | 0.0531 | 381 | 2.38 | 0.0177 | * | 0.0354 | * |
| Node | Fluid Intelligence | Head Motion | 0.0859 | 0.052 | 381 | 1.65 | 0.0994 | n.s. | 0.0994 | n.s. |
| Edge | Conscientiousness | None | 0.124 | 0.0506 | 382 | 2.45 | 0.0147 | * | 0.0173 | * |
| Edge | Externalizing | None | -0.157 | 0.0503 | 382 | -3.12 | 0.00194 | ** | 0.00582 | ** |
| Edge | Internalizing | None | -0.121 | 0.0506 | 382 | -2.39 | 0.0173 | * | 0.0173 | * |
| Edge | Conscientiousness | Demographics | 0.12 | 0.0509 | 380 | 2.35 | 0.0194 | * | 0.0291 | * |
| Edge | Externalizing | Demographics | -0.149 | 0.0506 | 380 | -2.94 | 0.00347 | ** | 0.01041 | * |
| Edge | Internalizing | Demographics | -0.108 | 0.0509 | 380 | -2.13 | 0.0342 | * | 0.0342 | * |
| Edge | Conscientiousness | Head Motion | 0.164 | 0.055 | 381 | 2.98 | 0.00307 | ** | 0.00921 | ** |
| Edge | Externalizing | Head Motion | -0.0969 | 0.0544 | 381 | -1.78 | 0.0759 | n.s. | 0.0759 | n.s. |
| Edge | Internalizing | Head Motion | -0.11 | 0.0552 | 381 | -1.99 | 0.0472 | * | 0.0708 | n.s. |
| Triangle | Externalizing | None | -0.136 | 0.0506 | 381 | -2.69 | 0.0074 | ** | 0.0148 | * |
| Triangle | Internalizing | None | -0.116 | 0.0506 | 381 | -2.3 | 0.0218 | * | 0.0218 | * |
| Triangle | Externalizing | Demographics | -0.14 | 0.0508 | 379 | -2.75 | 0.00628 | ** | 0.01256 | * |
| Triangle | Internalizing | Demographics | -0.107 | 0.0508 | 379 | -2.11 | 0.0355 | * | 0.0355 | * |
| Triangle | Externalizing | Head Motion | -0.0586 | 0.0571 | 380 | -1.03 | 0.305 | n.s. | 0.305 | n.s. |
| Triangle | Internalizing | Head Motion | -0.105 | 0.0576 | 380 | -1.82 | 0.069 | n.s. | 0.138 | n.s. |

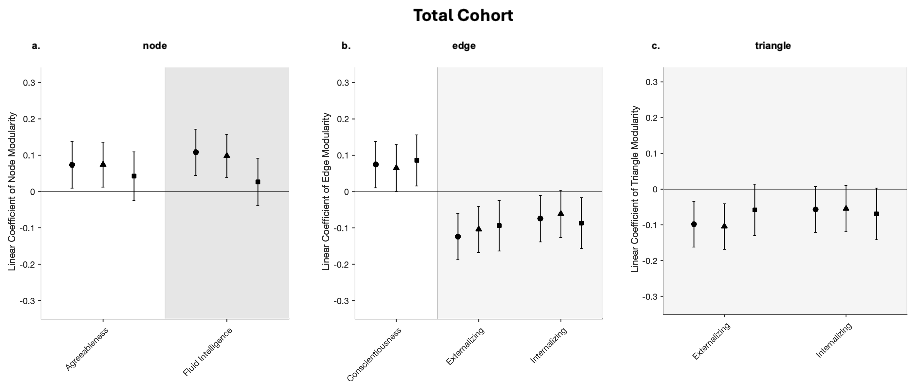

Fig. S12. 95% confidence intervals of the linear coefficients of the quality of modularity for the node, edge and triangle time series in the total cohort with demographics (age, sex, and family) and / or head motion controlled.

Table S12: Complete statistics of brain-behavior correlation in Fig. S12.

| **Simplex** | **Response Variable** | **Control Type** | **Estimate** | **Standard Error** | **DF** | **t-stat** | **p-value** | **Sig** | **p-val (BH corrected)** | **Sig (BH corrected)** |
| --- | --- | --- | --- | --- | --- | --- | --- | --- | --- | --- |
| Node | Agreeableness | None | 0.0734 | 0.0326 | 935 | 2.25 | 0.0246 | * | 0.0246 | * |
| Node | Fluid Intelligence | None | 0.108 | 0.0325 | 935 | 3.33 | 0.000895 | *** | 0.00179 | ** |
| Node | Agreeableness | Demographics | 0.0741 | 0.0315 | 933 | 2.35 | 0.0191 | * | 0.0191 | * |
| Node | Fluid Intelligence | Demographics | 0.0984 | 0.03 | 933 | 3.27 | 0.0011 | ** | 0.0022 | ** |
| Node | Agreeableness | Head Motion | 0.0428 | 0.0342 | 934 | 1.25 | 0.211 | n.s. | 0.422 | n.s. |
| Node | Fluid Intelligence | Head Motion | 0.0265 | 0.0331 | 934 | 0.801 | 0.423 | n.s. | 0.423 | n.s. |
| Node | Agreeableness | Head Motion + Family | 0.0421 | 0.0339 | 934 | 1.24 | 0.215 | n.s. | 0.262 | n.s. |
| Node | Fluid Intelligence | Head Motion + Family | 0.0353 | 0.0314 | 934 | 1.12 | 0.262 | n.s. | 0.262 | n.s. |
| Edge | Conscientiousness | None | 0.0745 | 0.0326 | 935 | 2.29 | 0.0224 | * | 0.0226 | * |
| Edge | Externalizing | None | -0.124 | 0.0324 | 935 | -3.82 | 0.000141 | *** | 0.000423 | *** |
| Edge | Internalizing | None | -0.0743 | 0.0326 | 935 | -2.28 | 0.0226 | * | 0.0226 | * |
| Edge | Conscientiousness | Demographics | 0.0651 | 0.0327 | 933 | 1.99 | 0.0464 | * | 0.0635 | n.s. |
| Edge | Externalizing | Demographics | -0.104 | 0.0326 | 933 | -3.19 | 0.00145 | ** | 0.00435 | ** |
| Edge | Internalizing | Demographics | -0.0614 | 0.0331 | 933 | -1.86 | 0.0635 | n.s. | 0.0635 | n.s. |
| Edge | Conscientiousness | Head Motion | 0.0858 | 0.0357 | 934 | 2.41 | 0.0164 | * | 0.0164 | * |
| Edge | Externalizing | Head Motion | -0.0938 | 0.0354 | 934 | -2.65 | 0.00825 | ** | 0.0164 | * |
| Edge | Internalizing | Head Motion | -0.0867 | 0.0357 | 934 | -2.43 | 0.0153 | * | 0.0164 | * |
| Edge | Conscientiousness | Head Motion + Family | 0.0807 | 0.0356 | 934 | 2.26 | 0.0238 | * | 0.0238 | * |
| Edge | Externalizing | Head Motion + Family | -0.0931 | 0.0357 | 934 | -2.61 | 0.00933 | ** | 0.0238 | * |
| Edge | Internalizing | Head Motion + Family | -0.0826 | 0.036 | 934 | -2.3 | 0.0219 | * | 0.0238 | * |
| Triangle | Externalizing | None | -0.098 | 0.0326 | 934 | -3.01 | 0.0027 | ** | 0.0054 | ** |
| Triangle | Internalizing | None | -0.057 | 0.0326 | 934 | -1.75 | 0.0809 | n.s. | 0.0809 | n.s. |
| Triangle | Externalizing | Demographics | -0.104 | 0.0324 | 932 | -3.22 | 0.00132 | ** | 0.00264 | ** |
| Triangle | Internalizing | Demographics | -0.0545 | 0.0329 | 932 | -1.65 | 0.0984 | n.s. | 0.0984 | n.s. |
| Triangle | Externalizing | Head Motion | -0.0581 | 0.0368 | 933 | -1.58 | 0.115 | n.s. | 0.115 | n.s. |
| Triangle | Internalizing | Head Motion | -0.0693 | 0.0369 | 933 | -1.88 | 0.061 | n.s. | 0.115 | n.s. |
| Triangle | Externalizing | Head Motion + Family | -0.0598 | 0.0371 | 933 | -1.61 | 0.107 | n.s. | 0.107 | n.s. |
| Triangle | Internalizing | Head Motion + Family | -0.0643 | 0.0372 | 933 | -1.73 | 0.0845 | n.s. | 0.107 | n.s. |

**S9 Stability of Edge Mapper Modularity across Sessions**

We next assessed whether the quality of modularity for edge time series Mapper graphs reflects a stable individual characteristic by testing its consistency across independent scanning sessions. We computed the correlation between the quality of modularity derived from the LR and RL scan sessions (Fig. S10). In the exploration cohort, the quality of modularity showed moderate inter-session reliability (r = 0.493, 95%-confidence interval: 0.413 – 0.565), significantly higher than that observed for node time series (r = 0.171, 95%-confidence interval: 0.0725 – 0.267, Steiger’s z = -4.94, p = 7.67E-7) . Triangle time series showed slightly lower reliability (r = 0.411, 95%-confidence interval: 0.324 – 0.491), although this difference was not statistically significant (Steiger’s z = 1.136, p = 0.256).

These findings replicated in the independent cohort, indicating that the quality of modularity for edge time series Mapper graphs captures a reproducible feature of individual brain dynamics rather than session-specific noise.

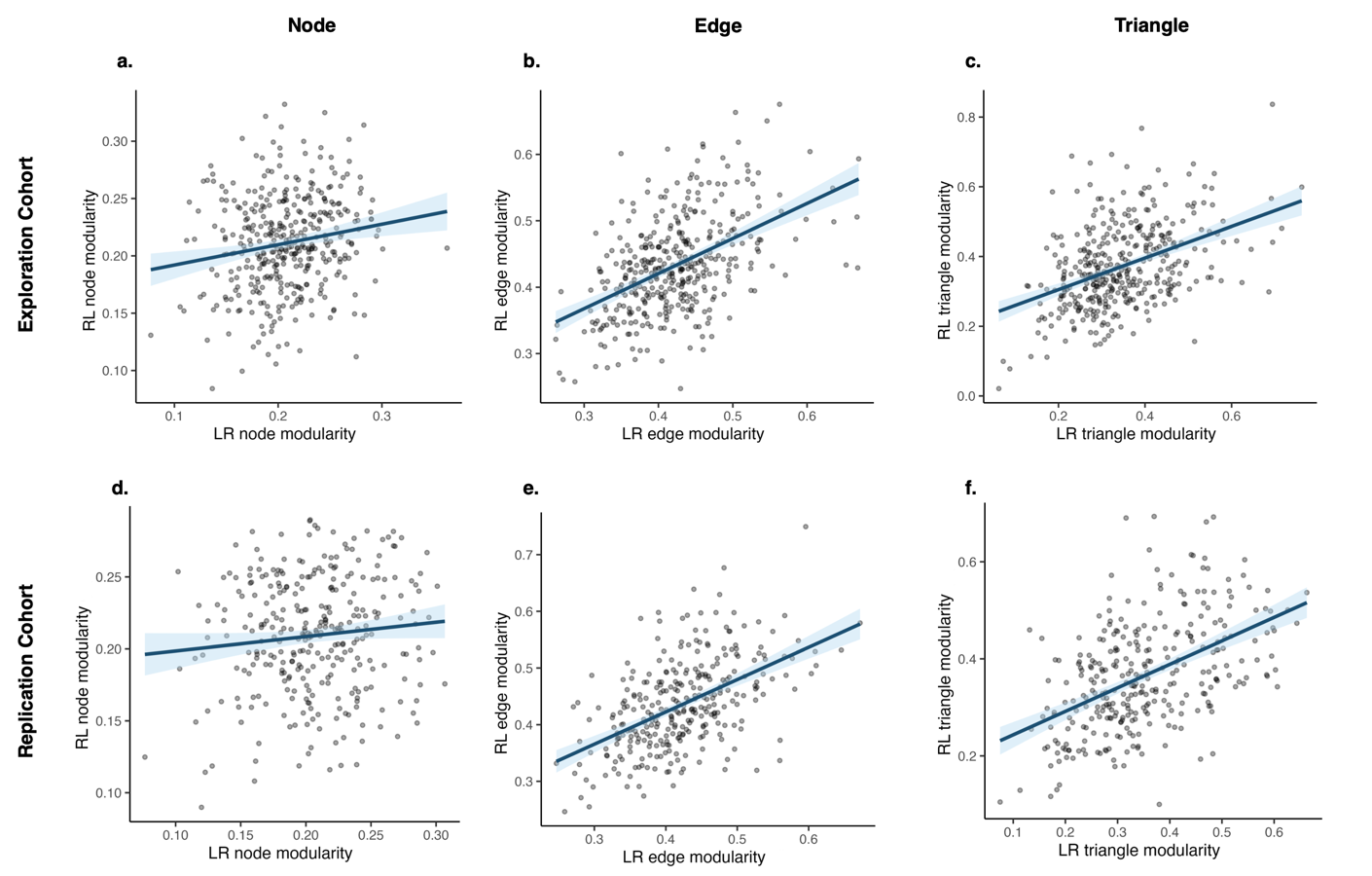
Fig. S13: Scatter plots of the quality of modularity of Mapper graphs for LR scan session and for RL scan sessions across subjects and across the choice of simplex (node time series, edge time series, and triangle time series).
